# Neural Pattern Change During Encoding of a Narrative Predicts Retrospective Duration Estimates

**DOI:** 10.1101/043075

**Authors:** Olga Lositsky, Janice Chen, Daniel Toker, Christopher J. Honey, Jordan L. Poppenk, Uri Hasson, Kenneth A. Norman

## Abstract

What mechanisms support our ability to estimate durations on the order of minutes? Behavioral studies in humans have shown that changes in contextual features lead to overestimation of past durations. Based on evidence that the medial temporal lobes and prefrontal cortex represent contextual features, we related the degree of fMRI pattern change in these regions with people’s subsequent duration estimates. After listening to a radio story in the scanner, participants were asked how much time had elapsed between pairs of clips from the story. Our ROI analysis found that the neural pattern distance between two clips at encoding was correlated with duration estimates in the right entorhinal cortex and right pars orbitalis. Moreover, a whole-brain searchlight analysis revealed a cluster spanning the right anterior temporal lobe. Our findings provide convergent support for the hypothesis that retrospective time judgments are driven by “drift” in contextual representations supported by these regions.

## Introduction

Imagine that you are at the bus stop when you run into a colleague and the two of you become engrossed in a conversation about memory research. After a few minutes, you realize that the bus still has not arrived. Before you look at your watch, you have some intuition for how long you’ve been waiting at the bus stop. Where does this intuition come from?

Estimation of durations lasting a few seconds has been probed in the neuroimaging, neuropsychology and neuropharmacology literatures (see Wittmann, 2013, for a review). On the other hand, the neural mechanisms underlying time perception on the scale of minutes have remained unexplored. This is particularly true of *retrospective* judgments, where individuals experience an interval without paying attention to time and must subsequently estimate the interval’s duration. In such cases, individuals must rely on information stored in memory to estimate duration. How is this accomplished?

Memory scholars have long posited that the same contextual cues that help us to retrieve an item from memory can also help us determine its recency. According to extant theories of context and memory (see Manning, Kahana, & Norman, 2014, for a review), *mental context* refers to aspects of our mental state that tend to persist over a relatively long time scale; this encompasses our representation of slowly-changing aspects of the external world (e.g., what room we are in) as well as other slowly-changing aspects of our internal mental state (e.g., our current plans). Crucially, these theories posit that slowly-changing contextual features can be episodically associated with more quickly-changing aspects of the world (e.g., stimuli that appear at a particular moment in time; Mensink & Raaijmakers, 1988; Howard & Kahana, 2002).

Bower (1972) first proposed that we could determine how long ago an item occurred by comparing our current context with the context associated with the remembered item. The similarity of these two context representations would reflect their temporal distance, with more similar representations associated with events that happened closer together in time. Thus, a slowly varying mental context could serve as a temporal tag (Polyn & Kahana, 2008). In parallel, researchers in the domain of retrospective time estimation have shown that the degree of context change is a better predictor of duration judgments than alternative explanations, such as the number of items remembered from the interval (Block & Reed, 1978; Block, 1990, 1992). Indeed, changes in task processing (Block & Reed, 1978; Sahakyan & Smith, 2014), environmental context (Block, 1982), and emotions (Pollatos, Laubrock, & Wittmann, 2014), as well as event boundaries (Poynter, 1983; Zakay, Tsal, Moses, & Shahar, 1994), lead to overestimation of past durations.

In our study, we set out to obtain neural evidence in support of the hypothesis that mental context change drives duration estimates. Specifically, we hypothesized that, in brain regions representing mental context, the degree of neural pattern change between two events (operationalized as change in multi-voxel patterns of fMRI activity) should predict participants’ estimates of how much time passed between those events.

Extensive prior work has implicated the medial temporal lobe (MTL) and lateral prefrontal cortex (PFC) in representing contextual information (Polyn & Kahana, 2008; for reviews of MTL contributions to representing context, see Eichenbaum, Yonelinas, & Ranganath, 2007, and Ritchey & Ranganath, 2012; for related computational modeling work, see Howard & Eichenbaum, 2013). In keeping with our hypothesis, multiple studies have obtained evidence linking neural pattern change in these regions to temporal memory judgments. Manns, Howard, & Eichenbaum (2007) recorded from rat hippocampus during an odor memory task; they found that greater change in hippocampal activity patterns between two stimuli predicted better memory for the order in which the stimuli occurred. In the human neuroimaging literature, Jenkins & Ranganath (2010) found that the degree to which activity patterns in rostrolateral prefrontal cortex changed during the encoding of a stimulus predicted better memory for the temporal position of that stimulus in the experiment. Jenkins & Ranganath (2016) also showed that greater pattern distance between two stimuli at encoding in the hippocampus, medial and anterior prefrontal cortex predicted better order memory. Only one study has directly related neural pattern drift to judgments of elapsed time in humans: Ezzyat & Davachi (2014) found that patterns of fMRI activity in left hippocampus were more similar for pairs of stimuli that were later estimated to have occurred closer together in time, despite equivalent time passage between all pairs (a little less than a minute).

While the Ezzyat & Davachi (2014) study provides support for our hypothesis, it has some limitations. First, in Ezzyat & Davachi (2014), participants estimated the temporal distance of stimuli that were linked to their contexts in an artificial way (by placing pictures of objects or famous faces on unrelated scene backgrounds); it is unclear whether these results will generalize to more naturalistic situations where events are linked through a narrative. Second, since participants performed the temporal memory test after each encoding run, they were not entirely naïve to the manipulation. Knowing that they would have to estimate durations between stimuli could have changed participants’ strategy and enhanced their attention to time (for evidence that estimating time prospectively engages different mechanisms, see Hicks, Miller, & Kinsbourne, 1976, and Zakay & Block, 2004). In the current study, we sought to address the above issues by eliciting temporal distance judgments for pairs of events that had occurred several minutes apart and that were embedded in the context of a rich naturalistic story; participants listened to the entire story before being informed about the temporal judgment task.

Based on the studies reviewed above, we predicted that neural pattern drift in medial temporal and lateral prefrontal regions might support duration estimation. In our study, we examined these regions of interest (ROIs), as well as a broader set of regions that have been implicated in fMRI studies of time estimation, including the inferior parietal cortex, putamen, insula and frontal operculum (see ***Box 1*** for a review). In addition to the ROI analysis, which examined activity patterns in masks that were anatomically defined, we performed a searchlight analysis, which examined activity patterns within small cubes over the whole brain.

Participants were scanned while they listened to a 25-minute science fiction radio story. Outside the scanner, they were surprised with a time perception test, in which they had to estimate how much time had passed between pairs of auditory clips from the story. Controlling for objective time, we found that the degree of neural pattern distance between two clips at the time of encoding predicted how much time an individual would later estimate passed between them. The effect was significant in two of our a priori ROIs – the right entorhinal cortex and the right pars orbitalis. Extending the anatomical analysis to all masks in cortex revealed an additional effect in the left caudal anterior cingulate cortex (ACC). Moreover, the whole-brain searchlight analysis yielded a significant cluster spanning the right anterior temporal lobe. Our results suggest that patterns of neural activity in these regions may carry contextual information that helps us make retrospective time judgments on the order of minutes.

#### Box 1. fMRI literature on prospective time estimation

As noted in the main text, only one study (Ezzyat & Davachi, 2014) has used fMRI to study retrospective estimation of time intervals lasting more than a few seconds. The vast majority of fMRI studies of time estimation have used prospective tasks, in which participants are asked to deliberately track the duration of a short stimulus or compare the duration of two stimuli. Such studies have repeatedly shown that activity in the putamen, insula, inferior frontal cortex (frontal operculum), and inferior parietal cortex increases as participants pay more attention to the duration of stimuli, as opposed to another time-varying attribute (Coull, Vidal, Nazarian, & Macar, 2004; Coull, 2004; Livesey, Wall, & Smith, 2007; Wiener, Turkeltaub, & Coslett, 2010; Wittmann, Simmons, Aron, & Paulus, 2010). Moreover, Dirnberger et al. (2012) showed that greater activity in the putamen and insula during encoding of aversive emotional pictures predicted better subsequent memory for those pictures, but only when their duration was overestimated relative to neutral images. This suggests that the putamen and insula might mediate the relationship between enhanced processing for emotional stimuli and subjective time dilation. Given the established role of these regions in time processing (albeit of a different sort) we included these regions in the set of a priori ROIs for our main fMRI analysis.

## Results

### Behavioral Results

***Figure 1*** shows the experimental design, which consisted of an fMRI session, followed immediately by a behavioral session. After listening to a 25-minute radio story in the scanner, participants were asked how much time had passed between 43 pairs of clips from the story. In actuality, 24 of the clip pairs had been presented 2 minutes apart in the story, while 19 of the clip pairs had been presented 6 minutes apart in the story (participants were not informed of this). Participants were able to estimate the duration of experienced minutes-long intervals far above chance, albeit with substantial intra- and inter-individual variability. On average, across participants, the 6-minute intervals (*M*=5.70 min, *SD*=3.06) were judged to be significantly longer than the 2-minute intervals (*M*=3.69 min, *SD*=1.96), *t*(17) = 5.20, *p* = 0.00007 (see ***Figure 2A***).

**Figure 1.**
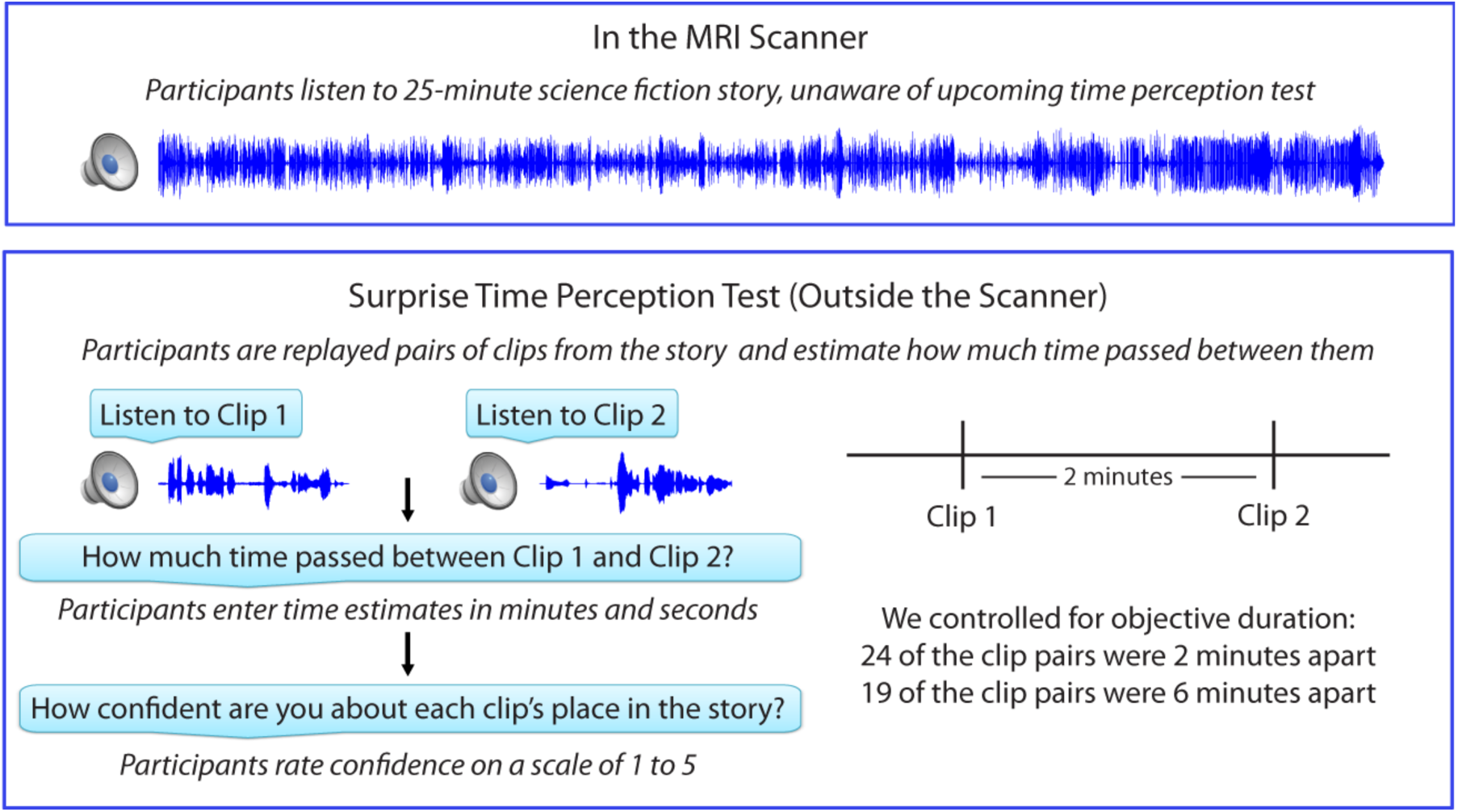
Experimental design.

**Figure 2.**
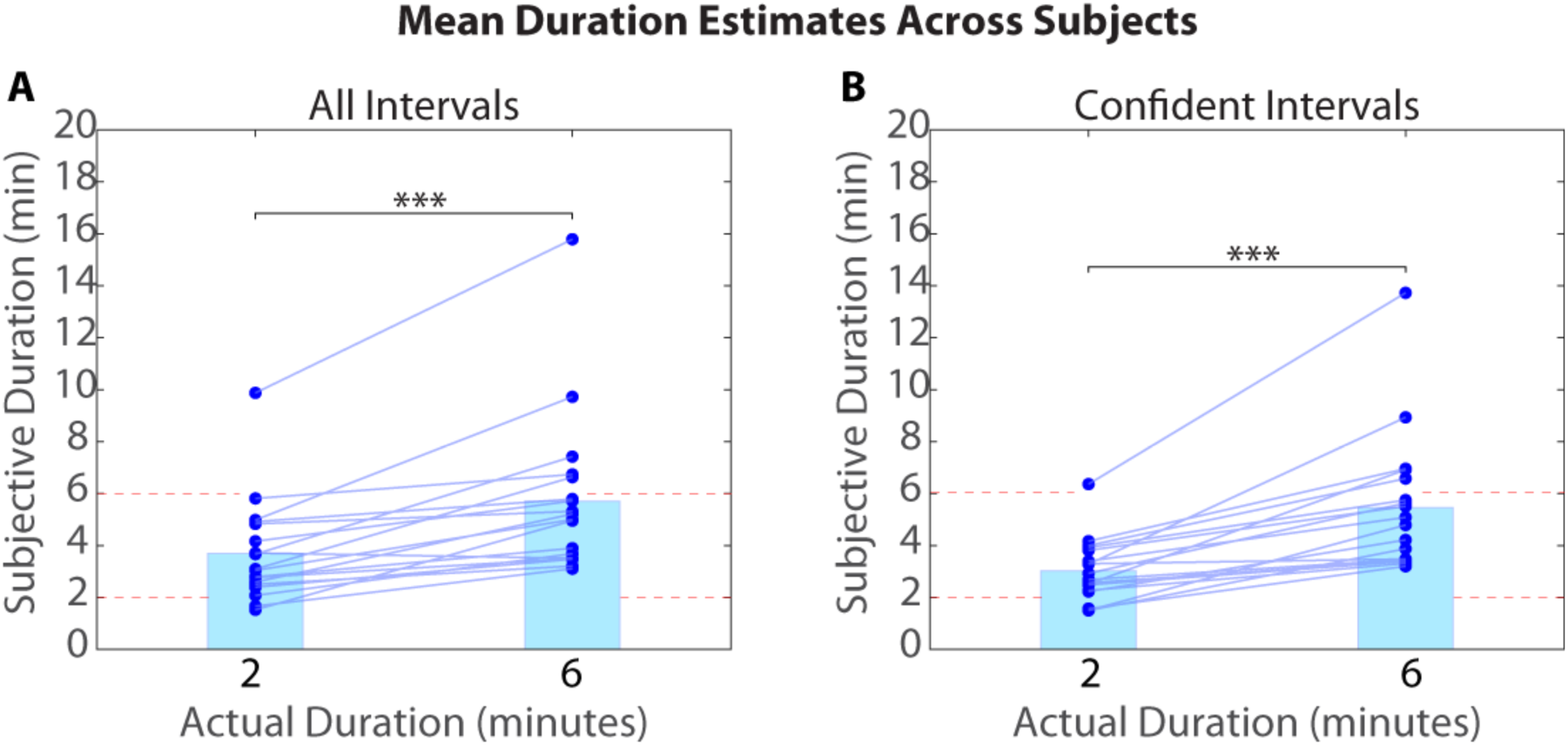
Mean duration estimates for all intervals (A) and confident intervals (B) as a function of their actual duration. Each blue circle represents the mean duration estimate for an individual participant within a given interval duration (2 or 6 minutes). The blue bar heights represent the global means for 2 and 6-minute intervals across intervals and participants.

As described in the Methods (see *Removing low-confidence intervals*), participants also provided confidence ratings reflecting their certainty about each clip’s place in the story. Based on this measure, we grouped each participant’s duration estimates into high-confidence and low-confidence intervals. To verify that participants were better at distinguishing 6-minute intervals from 2-minute intervals when they were confident, we calculated the difference between the mean duration estimates for 6-minute intervals and the mean duration estimates for 2-minute intervals for every participant. The difference score was significantly higher for high-confidence intervals (*M*=2.43, *SD*=1.82) than for all intervals (*M*=2.01, *SD*=1.64), *t*(17)=2.33, *p*=0.0324. Thus, participants were significantly more accurate at estimating an interval’s duration when they confidently remembered the temporal position of both clips delimiting that interval in the story (see ***Figure 2B***).

### Anatomical ROIs

We first tested whether pattern change in regions suggested by the literature to be important for representing temporal context (see *ROI Selection*) correlated with retrospective duration estimates. The analysis procedure is outlined in ***Figure 3***. As noted in the *Methods*, this analysis (as well as the searchlight analysis) was conducted only on high-confidence 2-minute intervals. 6-minute intervals were excluded from the fMRI analysis, since we could not successfully dissociate neural pattern change at this time scale from low-frequency scanner noise (see *Methodological challenges with analyzing pattern distance over long time scales* in the *Methods*).

**Figure 3.**
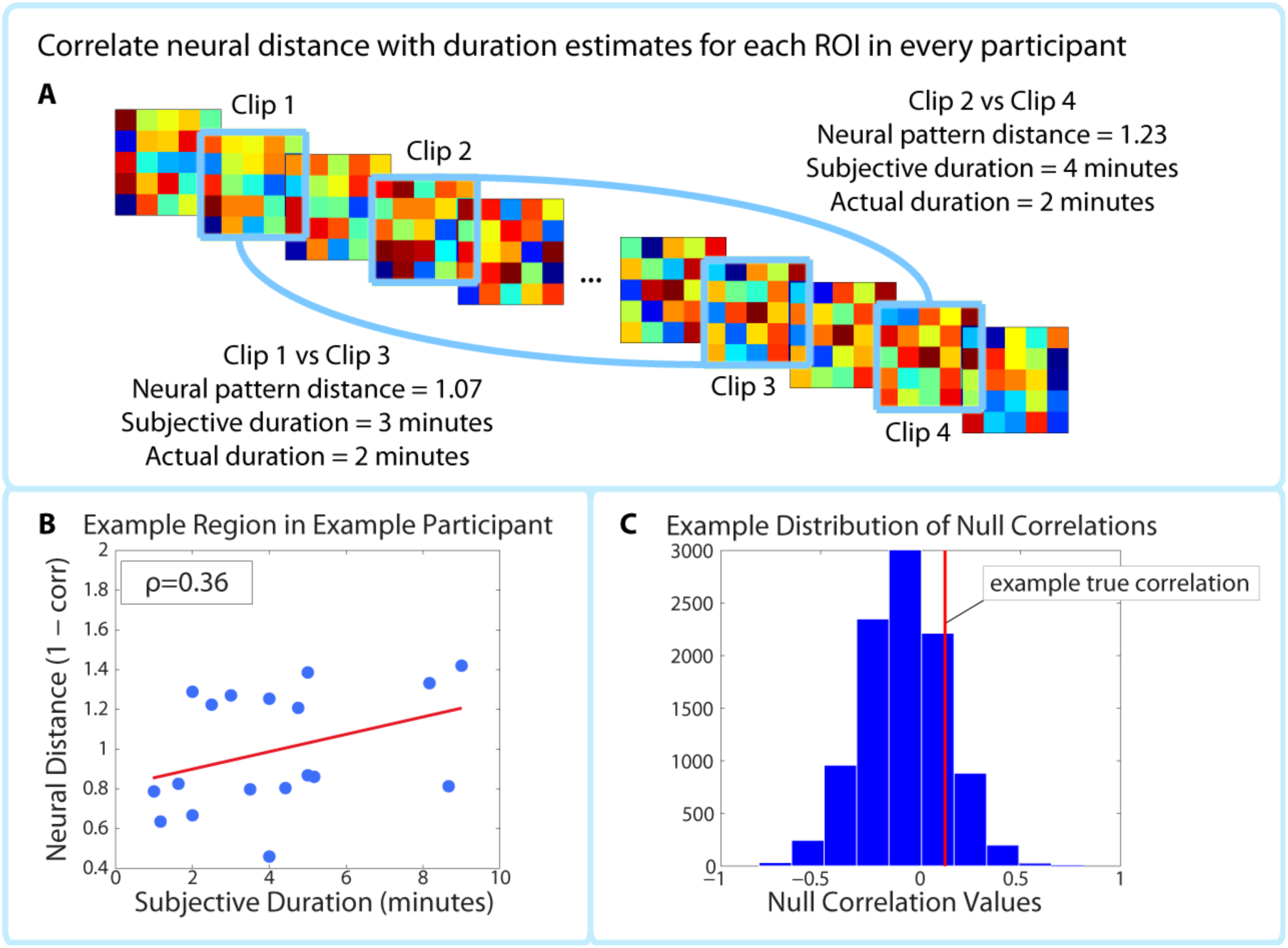
Correlating pattern distance with duration estimates within participants. For each ROI in each participant, the pattern distance between each pair of clips at encoding was correlated with the participant’s retrospective duration estimate (**A-B**). The top panel (**A**) shows two example intervals. The neural distance (1-Pearson’s *r*) between clips 2 and 4 (second interval) is greater than the neural distance between clips 1 and 3 (first interval), as is the subjective duration estimate. (**B**) shows the correlation between neural distance and duration estimates in a hypothetical region and participant. (**C**) We used a permutation test to generate 10,000 surrogate pattern distance vectors (see *Figure 3 - Supplement 1*), which we then used to obtain a distribution of null correlations between neural distances and duration estimates. For each ROI in each participant, we calculated the z-scored correlation value, which reflects the strength of the empirical correlation relative to the distribution of null correlations. For each ROI, we performed a random effects t-test to assess whether the z-score was reliably positive across participants. P-values from this t-test were then subjected to multiple comparisons correction.

Anatomical ROIs were derived from FreeSurfer cortical parcellation (Desikan et al., 2006) and from a probabilistic MTL atlas (Hindy & Turk-Browne, 2015). After calculating the empirical correlation between neural pattern distance and duration estimates in these ROIs (***Figure 3 A***), we used a phase randomization procedure (described in *Methods*) to obtain 10 000 null correlations for each ROI in every participant. This enabled us to calculate a Z-value for every ROI in every participant, which reflects the strength of the actual correlation between pattern distance and duration estimates relative to the distribution of null correlations (***Figure 3 C***). Here we report the regions whose Z-values were consistently positive across participants, corrected for multiple comparisons using False Discovery Rate.

Out of the regions selected a priori, the right entorhinal cortex and right pars orbitalis showed a significant positive correlation between pattern change and duration estimates for high-confidence 2-minute intervals (*q*<0.05). ***Figure 4*** shows the mean Z-values across participants for all a priori ROIs (16 in each hemisphere), including lateral prefrontal regions (top panel A), medial temporal lobe regions, insula, putamen, and inferior parietal cortex (bottom panel B). While a large number of these regions had Z-values that were positive across participants (e.g., left hippocampus, left entorhinal cortex, right perirhinal cortex, right amygdala, bilateral insula, and right caudal middle frontal cortex, *p*<0.05 uncorrected), we report only those that survived FDR correction.

**Figure 4.**
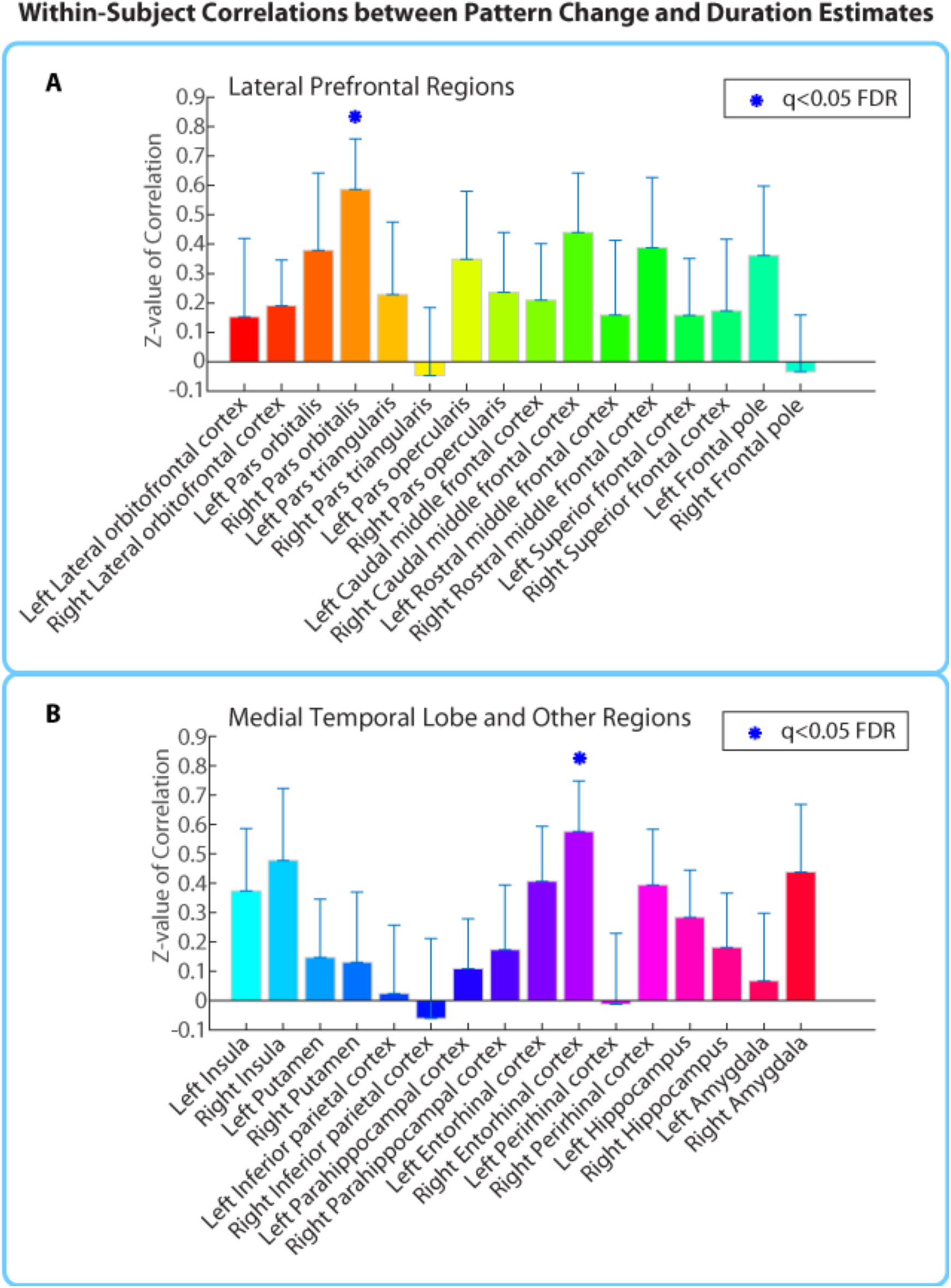
Mean Z-values (across all 18 participants) of correlations between pattern distance and duration estimates for the 16 a priori ROIs. Z-values were obtained from the phase randomization procedure and reflect the strength of the empirical correlation relative to the distribution of null correlations. Error bars represent standard errors of the mean. The blue dots over the right entorhinal cortex and right pars orbitalis indicate that these ROIs survived FDR correction at q<0.05.

As part of an exploratory search, we also performed this analysis on the other brain regions derived from FreeSurfer cortical parcellation. This included the 16 ROIs mentioned above, in addition to regions in the occipital lobe, parietal lobe, medial prefrontal cortex, lateral temporal lobe, basal ganglia, thalamus and brainstem (the complete list of regions can be found in ***Figure 4 – Supplement 1***). Out of the 84 regions tested (42 in each hemisphere), the right entorhinal cortex, right pars orbitalis, and left caudal anterior cingulate cortex (ACC) showed significant positive correlations between pattern change and duration estimates (*q*<0.1). This suggests that the right entorhinal cortex and right pars orbitalis, which were part of our list of a priori ROIs, contained effects that were apparent even after whole-brain correction, and reveals an additional effect in the left caudal ACC that we had not anticipated. ***Figure 4 – Supplement 2*** displays the locations of these three regions in MNI space.

### Whole-brain Searchlight

We also ran a cubic searchlight with 3×3×3 (27) voxels (972 mm^3^) through the entire brain, and tested for a correlation between pattern change and duration estimates in each searchlight. The same phase-randomization procedure that was used for the anatomical ROI analysis was also applied here; this procedure generates Z-values that reflect how likely we are to get this strong of a correlation by chance, given the frequency spectrum of the fMRI data. When excluding low-confidence intervals, we found a significant cluster in the right anterior temporal lobe (*p*=0.034, FWE-corrected; peak MNI coordinates (x, y, z) in mm: 45, ‐6, ‐21; cluster size=572 voxels in 3 mm MNI space). Small parts of the cluster also extended to the right posterior insula and right putamen (see ***Figure 5***).

**Figure 5.**
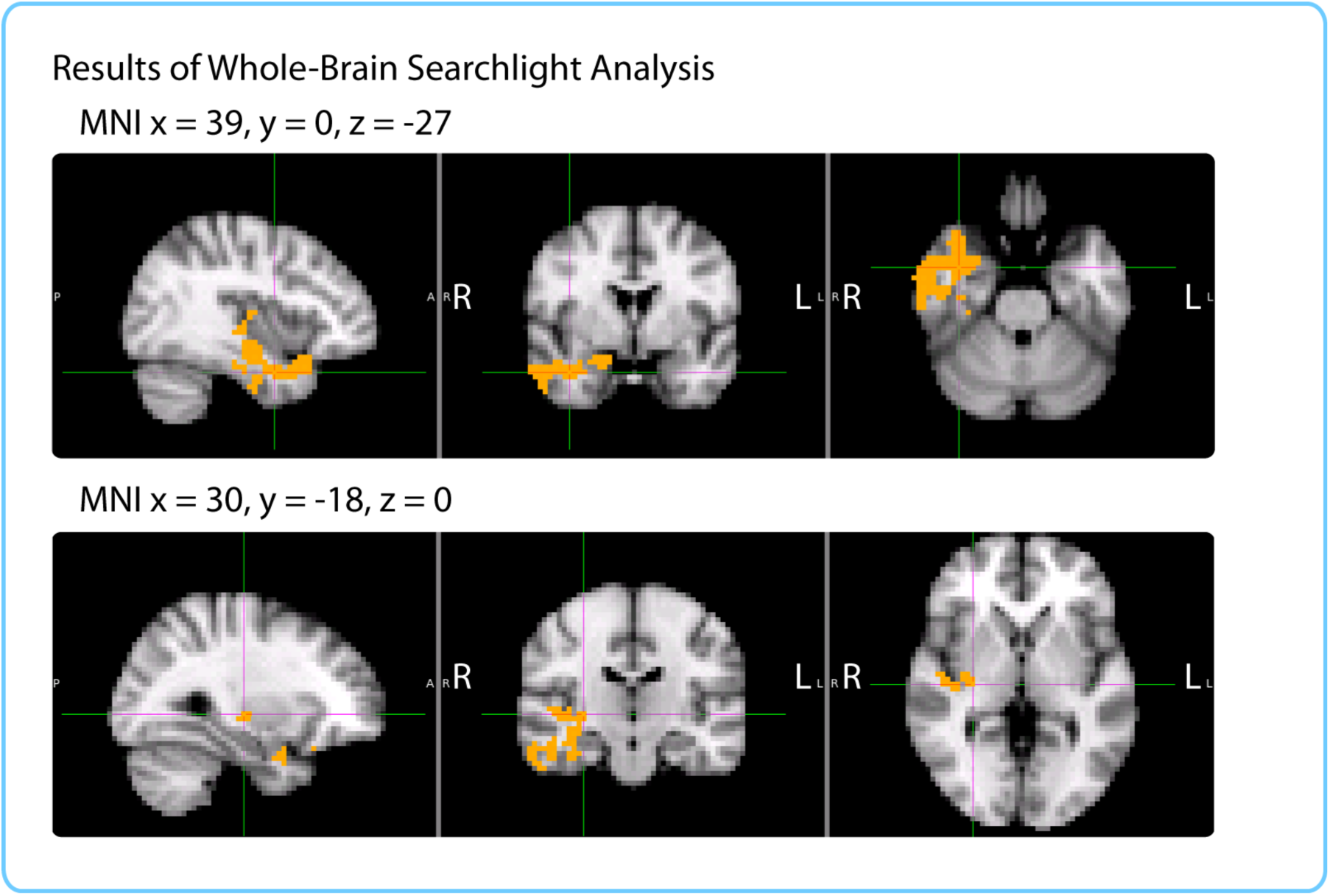
Results of a whole-brain 27-voxel cubic searchlight. Voxels in orange represent centers of searchlights that exhibited significant correlations between pattern change and duration estimates (p<0.05, FWE). The significant cluster had peak MNI coordinates (in mm): x=45, y= ‐6, z = ‐21.

### Comparing Results from ROI and Searchlight Analyses

The anatomical ROI and searchlight analyses revealed significant effects in somewhat different regions. For example, the ROI analysis yielded significant effects in the right pars orbitalis and left caudal ACC, while the searchlight did not; the significant searchlight cluster extended into the lateral temporal lobe and posterior insula, while the whole-brain ROI results did not. However, it is important to note that the regions we reported survived whole-brain multiple comparisons correction, and represent only the peaks of the effect. Looking at the unthresholded results reveals that both analyses converge in localizing the effect to a set of regions extending from the right anterior temporal lobe to the right inferior frontal cortex. ***Figure 6*** allows a comparison of the brain maps for both analyses with a more liberal threshold. To generate the brain map for the ROI analysis, we assigned the q-value of an ROI to each voxel in that ROI, and displayed all q-values < 0.3 (FDR). For the searchlight analysis, we displayed all clusters that were assigned p < 0.3 (FWE). This figure shows a sub-threshold searchlight cluster in the right inferior frontal cortex (panel B) that overlaps with the right pars orbitalis mask that was significant in the ROI analysis (panel A). Moreover, it shows a sub-threshold effect in the insula and superior temporal cortex ROIs (panel A), which overlap with the significant searchlight cluster (panel B).

**Figure 6.**
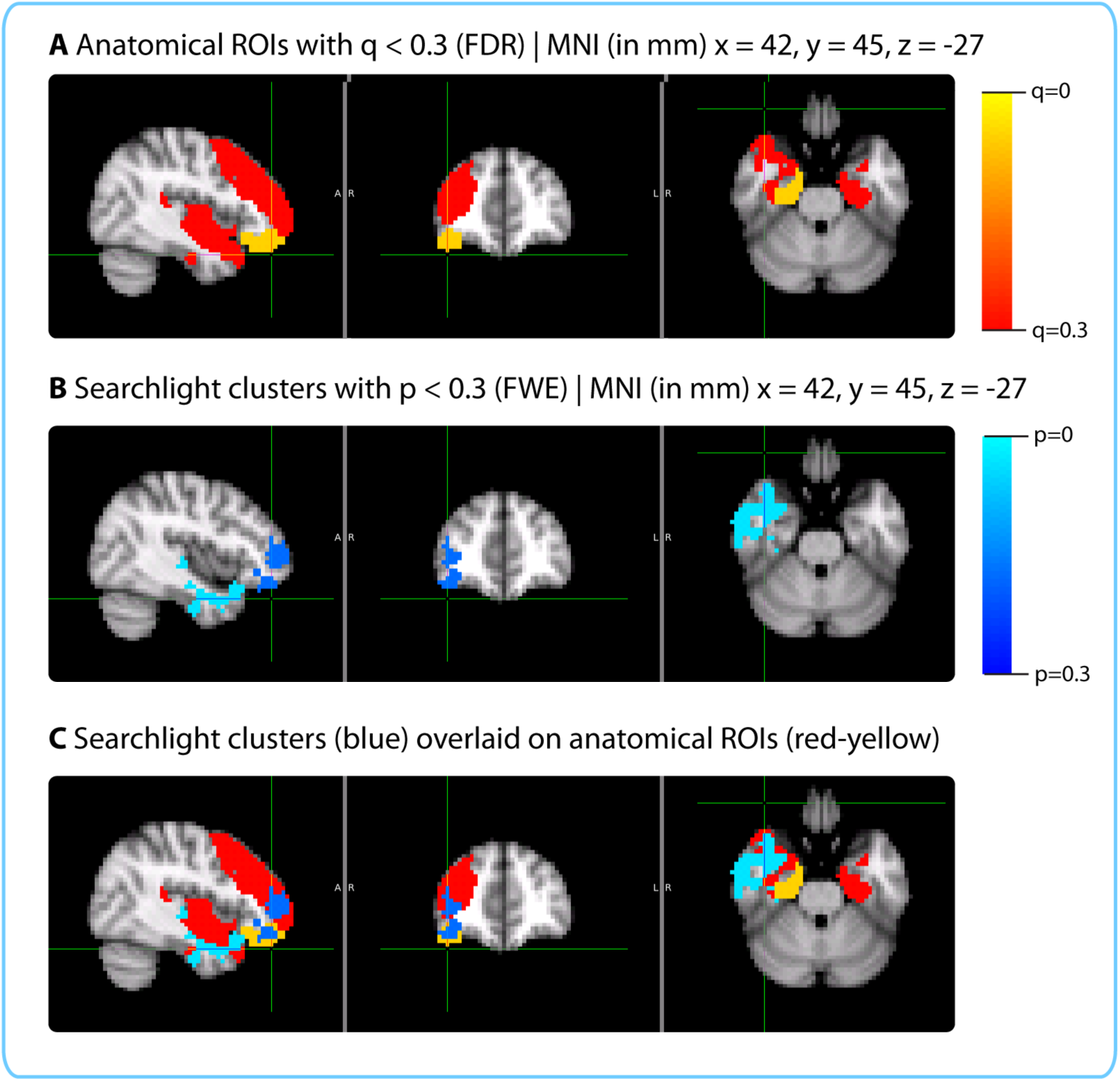
Comparison of the qualitative pattern of results revealed by the ROI and Searchlight analyses. Panel **A** shows the q-value map for the whole-brain ROI analysis, thresholded at *q*<0.3. Panel **B** shows searchlight clusters with *p*<0.3 (FWE). Panel **C** shows the searchlight clusters with *p*<0.3 (in dark and light blue) overlaid on top of the ROIs with *q*<0.3 (in red and yellow).

The difference in which regions passed the significance threshold is very likely due to the difference in shapes between the searchlight cube and the anatomical masks. For instance, the inferior and middle temporal cortex masks used in the ROI analysis encompass the entire inferior and middle temporal gyri and are too large to detect the multivariate effects in that region, which are confined to the anterior portion of the lateral temporal lobe. The small size of the searchlight cube enables it to better isolate this effect. On the other hand, the shapes of the pars orbitalis, entorhinal cortex and caudal ACC do not conform to a cubic searchlight, and are much better captured by the anatomical masks. Following the anatomy is particularly important for small, elongated regions like entorhinal cortex and caudal ACC, which are surrounded by white matter (our searchlight cubes could contain voxels in both grey and white matter, whereas the anatomical masks were limited to grey matter). Regions with such shapes are also less likely to be perfectly aligned across participants. For the searchlight analysis, individual participant searchlight maps needed to be transformed to MNI space in order to aggregate the results; consequently, imperfections in alignment can reduce the significance of searchlight results in these regions. On the other hand, the anatomical ROI analysis was performed entirely within participants, making it more suitable for idiosyncratically shaped regions.

### Factors Driving the Correlation between Pattern Change and Duration Estimates

We found that the neural pattern distance between two clips at encoding was correlated with participants’ subsequent temporal distance judgments in the right entorhinal cortex, right pars orbitalis, left caudal ACC and the right anterior temporal lobe. Our interpretation is that the above regions represent slowly varying contextual information and that duration estimates reflect the degree of mental context change between two clips. However, an alternative possibility is that participants judge the temporal distance between two clips purely based on the similarity between them (e.g. Are the same characters speaking? Is the background music the same? Is the topic of conversation similar?) If this is the case, participants do not need to retrieve the mental context associated with each clip and can estimate temporal distance based on the perceptual and semantic distance of the two clips at the time of the memory test. Thus, an alternative explanation of our neural findings is that patterns of activity in regions like the right entorhinal cortex and pars orbitalis reflect the moment-to-moment perceptual and semantic content of the story, but do not necessarily represent more abstract, slowly varying contextual features.

### Summary of control analyses

To rule out this possibility, we conducted two control behavioral studies. One group of participants indicated when event boundaries were occurring in the story. We show that the number of event boundaries between two clips correlates with duration estimates from our original participants, suggesting that their estimates were influenced by the content of the story in between two clips (rather than the similarity between the two clips alone.) A second group of participants was asked to complete the same time perception test without first listening to the story. Since these “naïve participants” had no memory of the story, they could *only* base their duration estimates on the similarity between the two clips. We show that duration estimates from these naïve participants do not correlate with the number of event boundaries between two clips, proving that the intervening content between clips does not influence duration estimates when participants have no memory of the story.

These behavioral controls provide evidence that our participants’ duration estimates were influenced by their memory of the story content in between two clips and could not be explained purely by the perceptual and semantic similarity between them. However, it is still possible that neural pattern change in the regions we found correlates with the component of duration estimates that is driven by perceptual and semantic content, rather than the component that is driven by slowly varying contextual information. To rule out this concern, we performed a within-interval (across participants) version of our main ROI analysis. This analysis holds constant the two clips whose pattern distance is being measured, and verifies whether individual differences in neural pattern distance for a given pair of clips correlate with individual differences in duration estimates for that interval. The within-interval analysis yielded the same constellation of regions in the right anterior temporal lobe and right inferior frontal cortex, including the right entorhinal cortex and right pars orbitalis, in addition to adjacent regions that had been sub-threshold in our main analysis. Further, we found that the effect sizes in the above two regions were similar for both analyses. If the neural pattern distance between two clips were driven by changes in clip content, we would have expected the effect to be larger for the across-interval, within-participants analysis (where story content differed across intervals) than for the across-participants, within-interval version of the analysis (where story content is held constant). The fact that the effect is similar in size for the two analyses suggests that the similarity in content between clips is not a major factor driving the observed correlation between neural pattern change and duration estimates.

Finally, we show that patterns in the right anterior temporal lobe and right inferior frontal regions we found change more gradually over time than patterns in most other brain regions, and that they change significantly more gradually than patterns in temporal lobe regions involved in auditory and language processing. Moreover, pattern change in the right entorhinal cortex correlates highly with pattern change in the right pars orbitalis, suggesting that the two regions may cooperate to represent different facets of a unified, slowly changing context signal.

### Correlation between number of event boundaries and duration estimates

First, we sought to replicate the finding that changes in contextual features would cause overestimation of durations in retrospect (see *Introduction*). We used the number of event boundaries between two clips in the story as a measure of the number of contextual changes, as event boundaries often encompass changes in scene, characters and conversation topic. A separate group of participants (n=9) listened to the story and was asked to press a button every time they felt an event boundary was occurring. These data were then averaged across participants to obtain the mean number of event boundaries inside each two-minute interval. We found that the mean number of boundaries in an interval was significantly correlated with the mean duration estimates from our original experiment (*r*=0.49, 95% CI [0.27, 0.57]; ***Figure 7***). This suggests that our participants’ retrospective duration estimates were influenced by the number of contextual changes that had occurred during an interval. However, it is possible that the number of event boundaries between two clips could influence the perceptual and semantic similarity of the clips themselves (e.g., clips from the same scene might be more similar than clips from different scenes). In that case, our participants’ duration estimates could correlate with the number of event boundaries, even if they are basing their estimates purely on the perceptual similarity between clips. To rule out this possibility, we tested whether the number of event boundaries would correlate with duration estimates from participants who could *only* judge temporal distance based on the similarity between clips, given that they had never heard the story.

**Figure 7.**
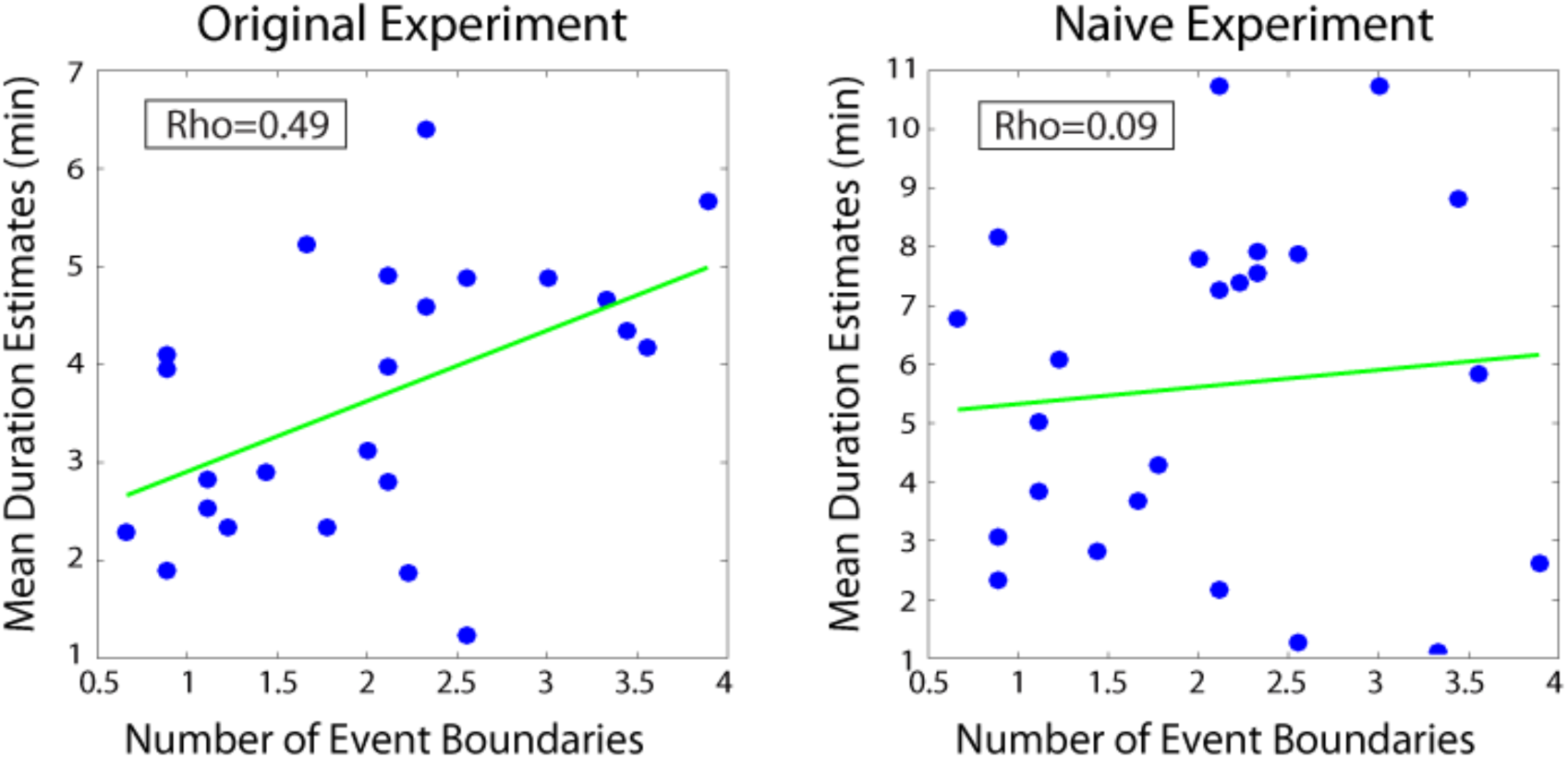
Mean duration estimates for 2-minute intervals as a function of the number of event boundaries in each interval. The number of event boundaries in an interval predicted retrospective duration estimates in our original experiment (left panel), but did not predict duration estimates of naïve participants (right panel) who had never heard the story. This suggests that the number of contextual changes between two clips influenced temporal distance judgments only when the content of the story between the two clips could be recalled.

### Naïve time perception test

An additional group of 17 participants who had never heard the story was administered an identical time perception test as our original participants. They were asked to try to estimate the amount of time that had elapsed between each pair of clips during the original telling of the story. Care was taken to ensure that participants understood the instructions, that they knew the story was 25 minutes long and that the maximum distance between two clips could not exceed that duration. During debriefing, participants reported making duration estimates based on the perceptual and semantic similarity between the two clips (e.g., which character voices were present, which background music was playing, the topic of conversation).

We found that naïve participants estimated 6-minute intervals (*M*=6.21 min, *SD*=1.91) to be longer than 2-minute intervals (*M*=5.63 min, *SD*=1.74; *t*(16)=2.62, *p*=0.019), suggesting that the similarity between two clips carried some information about the temporal distance between them. However, naïve participants were significantly less accurate at distinguishing 6-minute intervals from 2-minute intervals than our original participants who had heard the story. To quantify this, we calculated the difference between the mean duration estimates for 6-minute intervals and the mean duration estimates for 2-minute intervals for every participant (exactly as in the *Behavioral Results* section). The difference score was significantly higher for our original participants (*M*=2.01 min, *SD*=1.64 min) than for naïve participants (*M*=0.59 min, *SD*=0.91 min), *t*(26.86)= ‐3.22, *p*<0.005.

The inter-subject correlation in duration estimates was as strong for naïve participants (*M*=0.43, *SD*=0.18) as for our original participants (*M*=0A3, *SD*=0.25), suggesting that they used a consistent strategy to estimate durations. However, when we correlated duration estimates from our original group of participants with those of our naïve participants, we found that between-group correlations (*M*=0.18, *SD*=0.22) were significantly lower than the within-group correlations (*p*<0.0001, as assessed by a permutation test described in the *Methods*). This suggests that while both groups used a consistent strategy to estimate durations, the nature of the strategy differed across groups.

Most importantly, we found that the number of event boundaries in an interval did not significantly correlate with duration estimates of naïve participants (*r*=0.09, 95% CI [-0.05, 0.21]; ***Figure 7***). The correlation between the number of boundaries and duration estimates was significantly higher for our original participants than for naïve participants (*r*_*diff*_ = 0.40, 95% CI [0.15 0.56]).

These results suggest that duration estimates do not correlate with the number of contextual changes when participants are judging temporal distance based purely on the content of the clips. Rather, they support the interpretation that our original participants were recalling the content of the story in between two clips to estimate durations, and that contextual changes were a particularly strong driver of duration estimates.

However, it is still possible that pattern distance in the brain regions we found correlates with the component of duration estimates that is driven by the perceptual and semantic similarity between clips, rather than by contextual changes. To rule out this possibility, we performed a version of our main analysis that holds constant the perceptual and semantic similarity between two clips.

### Within-interval correlation between pattern distance and duration estimates

Our main ROI analysis was performed within participants and correlated the duration estimates for a given participant with that participant’s pattern distance vector across intervals. Results were then aggregated across participants. On the other hand, the “within-interval analysis” correlates duration estimates for a given interval across participants with the pattern distances for that interval (results are then aggregated across all 2-minute intervals). Rather than capturing variance within an individual across intervals of the story, this analysis captures variance across individuals for a given interval of the story. By performing the correlation for a given interval, we hold constant the perceptual and semantic content of the two clips and only leverage individual differences in how long the interval appeared retrospectively.

As described in the *Methods*, a permutation test was used to assess the statistical significance of each correlation. Duration estimates were scrambled across participants 10,000 times to obtain a distribution of null correlations, and Z-values were calculated for each interval, reflecting the strength of the empirical correlation relative to the distribution of null correlations. Finally, a right-tailed t-test was performed to assess whether the Z-values for a region were reliably above 0 across intervals. The *p*-values from this t-test were subjected to multiple comparisons correction using FDR.

Out of the regions selected *a priori*, the bilateral entorhinal cortex, right perirhinal cortex, right amygdala, right pars orbitalis, right lateral orbitofrontal cortex and bilateral insula showed a significant positive correlation between pattern change and duration estimates for 2-minute intervals (*q*<0.05). ***Figure 8*** shows the mean Z-values across intervals for all a priori ROIs (16 in each hemisphere).

**Figure 8.**
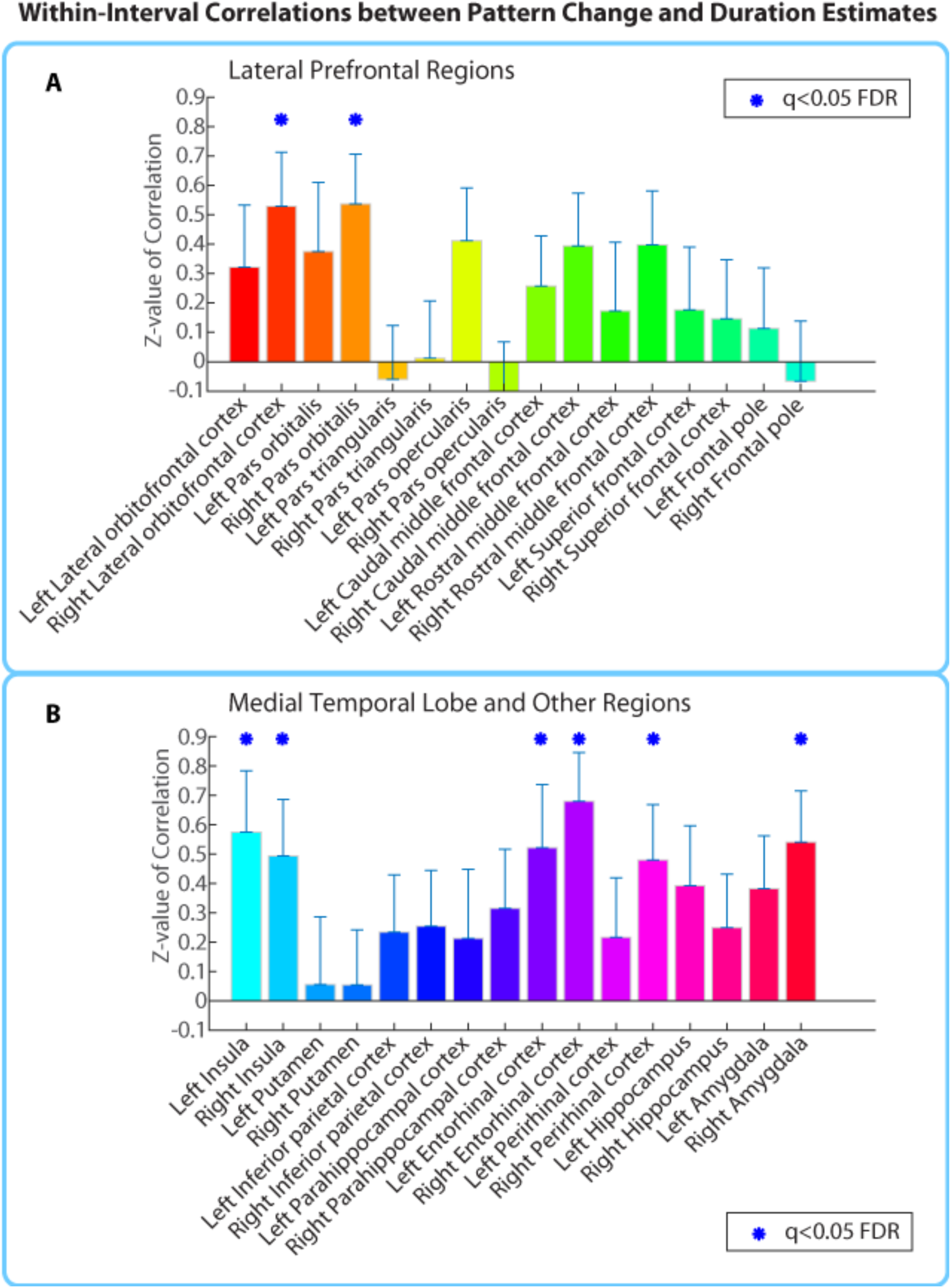
Mean Z-values (across all 2-minute intervals) of correlations between pattern distance and duration estimates for the 16 a priori ROIs. Error bars represent standard errors of the mean. Correlations between pattern change and duration estimates were performed across participants, separately for each interval.

Extending this analysis to the whole brain (same anatomical masks as in ***Figure 4 – Supplement 1***) revealed the same 8 ROIs listed above (suggesting all the effects were strong enough to survive whole-brain correction), in addition to the left caudal anterior cingulate cortex, and the right superior and inferior temporal cortices (*q*<0.1).

Note that the two regions revealed by our within-participant *a priori* ROI analysis, the right entorhinal cortex and right pars orbitalis, exhibited similar effect sizes in the within-interval analysis (Cohen’s *d* = 0.83 and 0.67, respectively) as in the within-participant analysis (Cohen’s *d* = 0.77 and 0.78, respectively).

If neural pattern distance in entorhinal cortex and pars orbitalis between two clips were driven by changes in clip content, we would have expected the correlation with duration estimates to be larger for the across-interval, within-participants analysis (where story content differed across intervals) than for the across-participants, within-interval version of the analysis (where story content is held constant). The fact that the effect sizes are similar, and that the qualitative pattern of brain regions matches that observed in the original analysis (the additional regions like perirhinal cortex, amygdala and insula had all been sub-threshold in the within-participant analysis) shows that differences in story content between two clips are not the main factor driving the correlation between duration estimates and neural pattern change.

### Patterns of activity in entorhinal cortex and pars orbitalis change slowly over time

To further probe the idea that the regions we found represent slowly changing contextual features, we assessed whether patterns of activity in these regions change more slowly over time than patterns of activity in regions known to be involved in auditory and language processing. To quantify the speed of pattern change, we obtained the mean auto-correlation function of the pattern in every region (see *Methods)*, and took the full-width half-maximum (FWHM) of this function as a measure of how slowly the pattern moves away from itself on average.

A right-tailed Wilcoxon signed-rank test indicated that the FWHMs in the right entorhinal cortex (*M*=18.9 TRs, *SD*=13.8 TRs) and right pars orbitalis (*M*=14.7 TRs, *SD*=8.0 TRs) were significantly larger across participants than the FWHMs in the right transverse temporal cortex (*M*=7.3 TRs, *SD*=1.2 TRs; *p*<0.00005 for the right entorhinal cortex and *p*<0.0005 for the right pars orbitalis), which encompasses primary auditory cortex (Destrieux, Fischl, Dale, & Halgren, 2010; Shapleske, Rossell, Woodruff, & David, 1999). The FWHMs in the entorhinal and pars orbitalis were also significantly larger than those in the right banks of the superior temporal sulcus (*M*=9.0 TRs, *SD*=2.1 TRs; *p*<0.001 for the right entorhinal cortex and *p*=0.0001 for the right pars orbitalis) and the right superior temporal cortex (*M*=11.0 TRs, *SD*=3.1 TRs; *p*<0.005 for the right entorhinal cortex and *p*<0.01 for the right pars orbitalis), regions involved in auditory processing and the early cortical stages of speech perception (Binder et al., 2000; Hickok & Poeppel, 2004). We also performed the above tests for the left caudal ACC, the region we found in our exploratory whole-brain analysis. The FWHMs in the left caudal ACC (*M*=8.3 TRs, *SD*=1.8 TRs) were significantly larger than those in the right transverse temporal cortex (*p*<0.01), but smaller than those in the right banks of the superior temporal sulcus (*p*=0.97) and right superior temporal cortex (*p*=1.0).

All of the p-values reported as significant also passed multiple comparisons correction with FDR (*q*<0.05).

We also ranked the mean FWHM in the regions we found relative to all the other masks in the brain (84 in total, 42 in each hemisphere). We found that the mean FWHM in the right entorhinal cortex (*M*=18.9) was the 3^rd^ largest in the entire brain, superseded only by the left temporal pole and the left medial orbitofrontal cortex. The mean FWHM in the right pars orbitalis (*M*=14.7) ranked 14^th^ in the brain and was superseded by the following regions: bilateral medial orbitofrontal cortex, bilateral temporal pole, bilateral entorhinal cortex, bilateral perirhinal cortex, left pars orbitalis, right inferior temporal cortex, right frontal pole and bilateral lateral orbitofrontal cortex.

These results show that the right entorhinal cortex and right pars orbitalis have some of the slowest pattern change in the entire brain, and the other slowest regions were adjacent regions of the orbitofrontal cortex and anterior temporal lobe, which we found in our searchlight analysis. On the other hand, the left caudal ACC did not have slower pattern change than most other brain regions and ranked 65^th^ out of 84 regions.

Since the masks used in our analysis were anatomically defined, they varied substantially in size. To ensure that differences in the speed of pattern change were not due to differences in ROI size (for instance, one could envision that patterns in smaller regions might change more gradually than patterns in large regions), we also performed all the above analyses after regressing out the vector of ROI sizes (number of voxels) out of the vector of FWHM values for each participant.

Regressing out ROI size did not affect any of the main findings above. The FWHMs in the right entorhinal cortex and right pars orbitalis were still significantly larger across participants than those in the right transverse temporal cortex (*p*<0.0005 for both regions) and those in the right banks of the superior temporal sulcus (*p*<0.001 and *p*<0.0005 respectively). However, they were no longer significantly larger than the FWHMs in the right superior temporal cortex (*p*=0.06 and *p*=0.09 respectively). Since the superior temporal cortex is a much larger ROI than entorhinal cortex or pars orbitalis, it is possible that regressing out ROI size obscures a true effect. The univariate analysis below addresses this issue and shows that individual voxels in superior temporal cortex exhibit faster signal change than voxels in entorhinal cortex or pars orbitalis. The FWHMs in the left caudal ACC were still significantly larger than those in the right transverse temporal cortex (*p*<0.005), but smaller than those in the right banks of the superior temporal sulcus (*p*=0.96) and right superior temporal cortex (*p*=1.0). All p-values reported as significant survived FDR correction at *q*<0.05.

Importantly, the ranking of the mean FWHMs in the right entorhinal cortex and right pars orbitalis relative to the rest of the brain was altered very slightly. The right entorhinal cortex now ranked 4^th^ in the brain, superseded only by the left and right medial orbitofrontal cortex and left temporal pole. The right pars orbitalis now ranked 10^th^ in the brain and was superseded only by the bilateral medial orbitofrontal cortex, bilateral temporal pole, right entorhinal cortex, right inferior temporal cortex, bilateral lateral orbitofrontal cortex and left perirhinal cortex. Again, the left caudal ACC did not exhibit slower pattern change than most other regions and ranked 66^th^ out of 84 regions.

These results supported our interpretation that patterns in the right entorhinal cortex and pars orbitalis change more slowly than in most other regions because they process abstract information that changes slowly over time (rather than an artifact of ROI size). However, it was still possible that regressing out ROI size would only remove a linear relationship between ROI size and the speed of pattern change but could not rule out a non-linear relationship. To address this concern more thoroughly, we performed the above analysis for every voxel individually. Rather than calculating the mean autocorrelation function of the pattern in every region, we calculated the auto-correlation function of every voxel’s time course and averaged the auto-correlation functions across all the voxels in a given region. The FWHM was then computed for this mean autocorrelation derived from individual voxel time courses.

This univariate analysis yielded very similar results qualitatively. The FWHMs in the right entorhinal cortex (*M*=23 TRs, *SD*=15.6 TRs) and right pars orbitalis (*M*=17.1 TRs, *SD*=7.7 TRs) were significantly larger across participants than those in the right transverse temporal cortex (*M*=7.9 TRs, *SD*=1.2 TRs; *p*<0.00005 and *p*<0.0005 respectively), the right banks of the superior temporal sulcus (*M*=8.8 TRs, *SD*=1.7 TRs; *p*<0.0005 and *p*<0.00005 respectively) and the right superior temporal cortex (*M*=10.3 TRs, *SD*=2.4 TRs; *p*<0.0005 and *p*<0.0001 respectively). However, the FWHMs in the left caudal ACC (*M*=9.22 TRs; *SD*=3.8 TRs) were no longer significantly larger than those in the right transverse temporal cortex (*p*=0.059), the right banks of the superior temporal sulcus (*p*=0.42) or the right superior temporal cortex (*p*=0.98). In terms of ranking, the right entorhinal cortex now ranked 1^st^, displaying the largest mean FWHM value in the entire brain. The right pars orbitalis now ranked 11^th^ and was superseded by the same regions listed above, including the bilateral entorhinal cortex, bilateral frontal pole, bilateral temporal pole, bilateral perirhinal cortex, bilateral medial orbitofrontal cortex and the left pars orbitalis. On the other hand, the left caudal ACC ranked 45^th^ out of 84 brain regions, suggesting that its patterns did not change more slowly than most other regions.

Taken together, all three variants of the analysis show that the right entorhinal and right pars orbitalis, along with neighboring regions of the anterior and medial temporal lobe, orbitofrontal cortex and frontal pole, have the slowest pattern change in the brain. These results do not seem to be due to differences in the sizes of the anatomical masks and suggest that the regions found in our ROI analysis process information that changes slowly over time. Our findings are consistent with those of Stephens, Honey, & Hasson (2013), who showed that auditory cortex regions processing momentary stimulus features had intrinsically faster dynamics than higher-order regions that integrated information over longer time scales (see also Lerner, Honey, Silbert, & Hasson, 2011).

### Pattern distances in the right entorhinal cortex and right pars orbitalis are correlated

While we did not predict that all regions whose pattern distance predicted duration estimates should represent the same contextual information, we hypothesized that these regions would have correlated pattern change if they are part of a network that integrates the various features comprising mental context. To explore this, we first averaged the pattern distance values across participants for each of the 24 pairs of clips (which were 2 minutes apart). This step was performed based on findings that averaging across participants helps to reduce correlations within brains that are driven by respiration, heart rate and head motion, and helps to isolate the correlations across regions that are driven by neural processing of the story (Simony et al., in press). The correlation between the mean pattern distance vectors in the right entorhinal cortex and right pars orbitalis was *r* = 0.73. In order to interpret the magnitude of this correlation, we also calculated the correlation between every possible pair of mean pattern distance vectors (for all 84 anatomical masks). This resulted in a distribution of 3486 correlations – one for every possible pair of regions. Out of 3486 pairs of regions, only 242 exhibited a correlation that was higher than the one observed between the right entorhinal and right pars orbitalis. Thus, the correlation between the pattern distances in these two regions is higher than for 93% of region pairs.

To ascertain that the correlation between these regions was not spuriously high because of the auto-correlation in the pattern distance signal, we performed a phase randomization procedure (see *Methods*) to assess the likelihood of this correlation magnitude, given the frequency spectra of the pattern distance vectors. We generated 1000 surrogate pattern distance vectors for each region, and correlated each surrogate entorhinal vector with each surrogate pars orbitalis vector, resulting in a distribution of 1,000,000 null correlations. The likelihood of obtaining a correlation of *r*=0.73 or higher by chance was *p*=0.0011.

We also verified whether the pattern dissimilarity values in the right entorhinal cortex and right pars orbitalis were correlated with those in the left caudal anterior cingulate cortex (ACC), the region yielded by extending our anatomical ROI analysis to the whole brain. The pattern change in the left caudal ACC was substantially less correlated with pattern change in the right entorhinal cortex (*r*=0.23, higher than only 14.7% of region pairs; *p*=0.19 based on the phase randomization procedure) and the right pars orbitalis (*r*=0.44, higher than 41.1% of region pairs; *p*=0.051 based on the phase randomization procedure).

Together with the finding that patterns in the left caudal ACC change more rapidly than those in the entorhinal cortex or pars orbitalis, this suggests that pattern change in the ACC captures a qualitatively different signal than the one represented in the entorhinal cortex and pars orbitalis.

### Story position effects can not explain the correlation between duration estimates and neural pattern change

We found that duration estimates systematically decreased as a function of position in the story, with earlier intervals being estimated as longer than later intervals (***Figure 9A***). The correlation between the estimated duration of an interval and its time in the story was consistently negative across participants (*M*= ‐0.40, *SD*= 0.22; *t*(16)= ‐7.59, *p*<0.00001; we defined the time of the interval in the story as the middle time point of each 2-minute interval, half-way between the two clips delimiting it). If neural pattern change also decreased as a function of position in the story, it might be possible to explain the observed correlation between duration estimates and pattern change in terms of both measures being correlated with story position.

**Figure 9.**
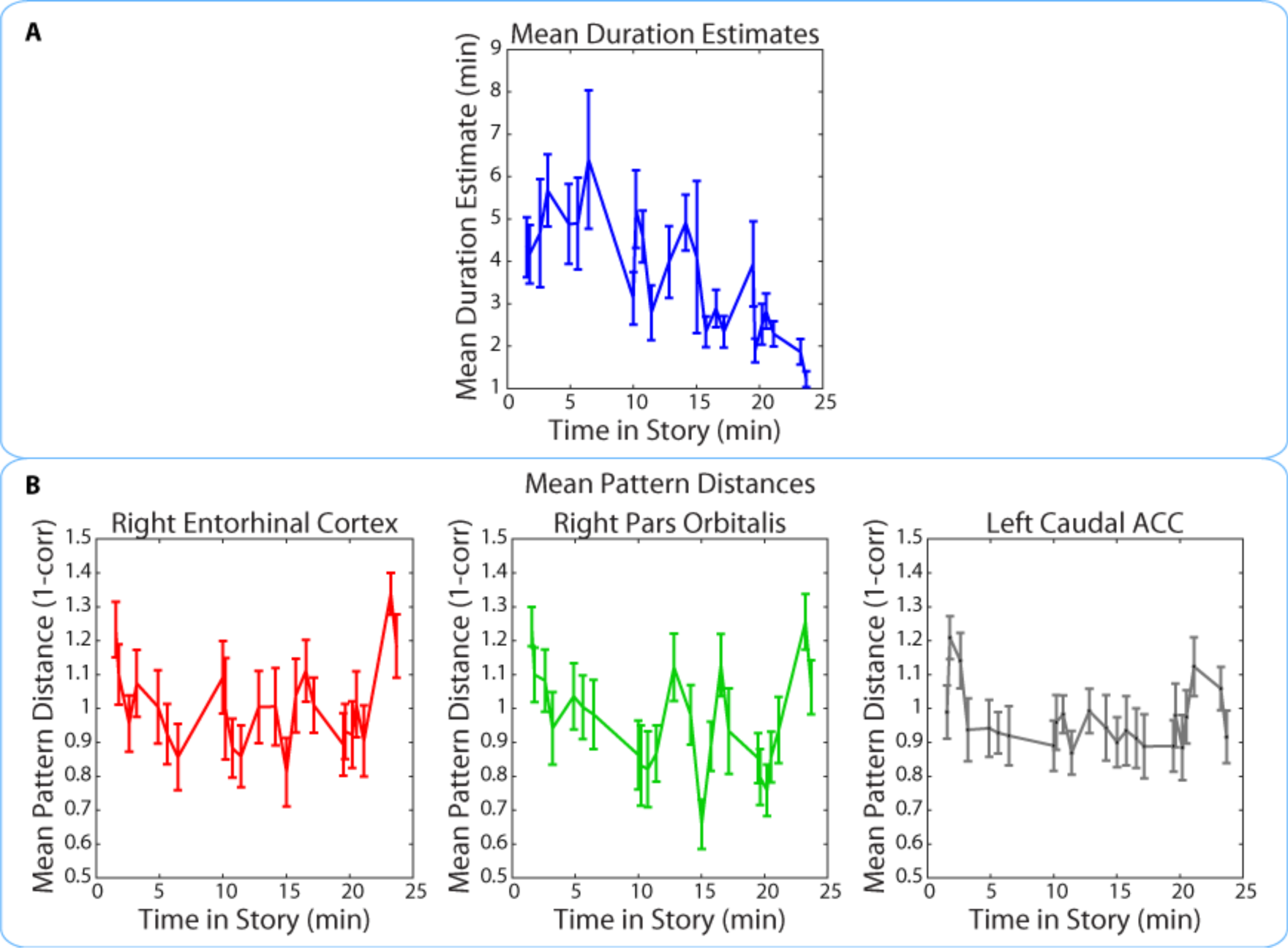
Mean duration estimates and pattern distances (across participants) for all 2-minute intervals as a function of the interval’s position in the story. The middle time point of each 2-minute interval (half-way between the two clips delimiting it) was chosen as the x-coordinate.

Importantly, the pattern dissimilarity values in right entorhinal cortex and right pars orbitalis did not exhibit the same overall decrease across time. In fact, there was no consistent correlation between pattern change during an interval and its time in the story for the right entorhinal cortex (*M*=0.03, *SD*=0.21; *t*(16)= 0.65; *p*=0.53) or the right pars orbitalis (*M*=-0.10, *SD*=0.22; *t*(16)= ‐1.83, *p*=0.09). This correlation was also not reliable for the left caudal ACC (*M*=-0.05, *SD*=0.18; *t*(16)=-1.15, *p*=0.27), the region found in our exploratory whole-brain anatomical analysis. These results suggest that the relationship between duration estimates and pattern dissimilarity in these regions was not driven by a shared effect of story position. Rather, it seems that pattern dissimilarity in these regions correlated with more fine-grained variations in the estimated durations of nearby intervals (***Figure 9B***).

## Discussion

While human and animal time perception has been a subject of intense empirical investigation (see Wittmann, 2013), most neuroimaging studies have tested its mechanisms on the scale of milliseconds to seconds and neglected paradigms in which long-term memory plays an important role. Such studies have typically employed *prospective paradigms*, in which participants must deliberately attend to the duration of a stimulus. However, behavioral studies in humans have consistently demonstrated that *retrospective paradigms*, in which participants are asked to estimate the duration of an elapsed interval from memory, tap into different cognitive mechanisms from prospective ones (Hicks et al., 1976; Zakay & Block, 2004; Block & Zakay, 2008). In retrospective paradigms, changes in spatial, emotional and cognitive context tend to modulate estimates of elapsed time (Block & Reed, 1978; Block, 1992; Sahakyan & Smith, 2014; Pollatos et al., 2014).

In the present study, we used changes in patterns of BOLD activity as a proxy for mental context change. We sought to extend previous neuroimaging work by testing whether neural pattern change predicts duration estimates on the scale of several minutes and in a more naturalistic setting, where spatial location, situational inference, characters, and emotional elements can all drive contextual change.

Participants were scanned while they listened to a 25-minute radio story and were subsequently asked how much time (in minutes and seconds) had elapsed between pairs of clips from the story (all pairs were in fact two minutes apart). Using this approach, we were able to probe retrospective duration memory repeatedly within participants without needing to interrupt the encoding of the story. This allowed us to leverage within-participant variability in neural pattern change and relate it to a participant’s retrospective duration estimates.

Using a within-participant anatomical ROI analysis (encompassing 16 regions selected a priori), we found that neural pattern distance in the right entorhinal cortex and right pars orbitalis at the time of encoding was correlated with subsequent duration estimates. Extending this analysis to all anatomical ROIs in cortex revealed an additional effect in the left caudal anterior cingulate cortex (ACC). These results converged qualitatively with the results of our whole-brain searchlight analysis, which revealed a significant cluster spanning the right anterior temporal lobe and extended to a sub-threshold cluster in the right inferior frontal cortex.

To test our interpretation that both duration estimates and neural pattern distance were driven by contextual change, we asked a separate group of participants to identify event boundaries in the story. We found that the number of event boundaries between two clips was very highly correlated with participants’ subsequent duration estimates. Importantly, the number of event boundaries did not predict duration estimates for a separate group of “naïve” participants, who had been asked to estimate the elapsed time between clips without first hearing the story. This showed that simply hearing the content of the two clips was not sufficient to infer the number of contextual boundaries between them. These behavioral experiments provide evidence that retrospective duration estimates were indeed influenced by memory for intervening contextual changes between clips.

In addition, we sought to rule out the possibility that neural pattern distance between two clips reflected only the perceptual or semantic similarity between them, rather than the degree of mental context change. We performed a within-interval anatomical ROI analysis, in which pattern distances for the same pair of clips were correlated with duration estimates *across participants*. This analysis yielded effects of the same size in the right entorhinal cortex, right pars orbitalis and left caudal ACC, as well as an additional set of regions like the right lateral orbitofrontal cortex, left entorhinal, right perirhinal cortex, right amygdala and bilateral insula. In other words, pattern distance in the regions we found predicted variability in duration estimates even when the perceptual and semantic distance of the clips was controlled as much as possible, suggesting that their signal may capture idiosyncratic differences in mental context that cannot be predicted from the stimulus alone.

Finally, measuring the speed of neural pattern change in all anatomical ROIs revealed that the right entorhinal cortex, right pars orbitalis, as well as adjacent regions of the MTL, temporal pole and orbitofrontal cortex, had some of the slowest changing patterns in the entire brain. This is in line with findings that brain regions at the top of the processing hierarchy (furthest from the primary perceptual areas) integrate information over longer time scales and are therefore best suited for representing abstract information extracted from multiple streams of sensory observations (Stephens, Honey, & Hasson, 2013; Lerner et al., 2011). We also found that the right pars orbitalis and right entorhinal cortex had some of the most correlated pattern change out of any pair of brain regions, making them strong candidates for integrating various features that comprise mental context into a unified representation that changes slowly over time.

Below, we discuss how the medial temporal, lateral prefrontal and anterior cingulate regions we found most consistently throughout all our analyses relate to previous findings in the literature.

Multiple lines of evidence have suggested an important role for the entorhinal cortex in representing relationships between the spatial environment, task and incoming stimuli. Lesions of the lateral entorhinal cortex in rodents have shown that this region is necessary for discriminating between novel and familiar associations of object and place, object and non-spatial context, or place and context, while leaving non-associative forms of memory unaffected (Buckmaster, Eichenbaum, Amaral, Suzuki, & Rapp, 2004; Wilson, Watanabe, Milner, & Ainge, 2013; Wilson, Langston, et al., 2013). Moreover, electrophysiological recordings in rats performing a spatial memory task showed that neurons in the medial entorhinal cortex exhibited greater context sensitivity and greater modulation by task-relevant mnemonic information than hippocampal neurons, while hippocampal neurons carried more specific spatial information (Lipton, White, & Eichenbaum, 2007). Medial entorhinal neurons also exhibited longer firing periods, which led the authors to propose that they could bind a series of hippocampal representations of distinct events (Lipton & Eichenbaum, 2008). Thus, changes in distributed entorhinal activity patterns on the scale of minutes might represent changes in contextual elements that are later retrieved to make duration judgments (for theoretical discussion of the role of entorhinal cortex in contextual representation, see Howard, Fotedar, Datey, & Hasselmo, 2005).

While the right entorhinal cortex was the only medial temporal lobe region that survived FDR correction in both our within-participant and within-interval ROI analyses, our whole-brain searchlight found a significant relationship between pattern change and duration estimates in an extensive cluster that overlapped with a few voxels in the right hippocampus, the right perirhinal cortex, and right temporal pole. Moreover, our within-interval ROI analysis yielded significant effects in the bilateral entorhinal cortex, right perirhinal and right amygdala.

Two previous studies, Noulhiane et al. (2007) and Ezzyat and Davachi (2014), have directly implicated the MTL in retrospective time estimation in humans. Ezzyat and Davachi (2014) scanned participants while they were presented with trial-unique faces and objects on a scene background, which changed every four trials. After each run, participants were asked whether pairs of stimuli had occurred close together or far apart in time (all pairs were about 50 seconds apart). They found that neural pattern distance in the left hippocampus at the time of encoding was greater for pairs of stimuli later rated as “far apart”, though only when the stimuli were separated by a scene change. Noulhiane et al. (2007) used a retrospective behavioral paradigm similar to ours in patients with unilateral MTL lesions. In that study, participants were asked to estimate the temporal distance between object pictures that had been randomly inserted into a silent documentary film. They found that the degree of left entorhinal, left perirhinal and left temporopolar cortex damage correlated with the degree to which patients overestimated minutes-long intervals in retrospect. (For related evidence from the animal literature, see Jacobs, Allen, Nguyen, & Fortin, 2013, who showed that bilateral inactivation of the hippocampus impaired rats’ ability to discriminate between similarly long durations, like 8 and 12 minutes, but not between less similar intervals, like 3 and 12 minutes.)

Our ROI and searchlight results are in line with the above set of findings, and suggest that patients with anterior MTL lesions might be impaired in retrospective time estimation because patterns of activity in entorhinal, perirhinal, and temporopolar cortex encode contextual changes on the scale of minutes. The set of regions we found is more extensive than those in Ezzyat & Davachi (2014) and mostly right-lateralized. It is possible that the difference in the extent of our effects could be explained by the difference in paradigm. In both the Noulhiane (2007) and Ezzyat & Davachi (2014) studies, the links between objects and their context had to be deliberately constructed. In our study, the clips whose temporal distance participants estimated were excerpts from a story, and therefore strongly linked with a situational, spatial, and emotional context. Thus, it is possible that activity patterns in a more extensive cluster tracked temporal distance estimates because our auditory story caused changes in a broader set of contextual features.

Beyond the medial temporal lobe, our ROI analysis revealed a relationship between pattern distance and duration estimates in the right pars orbitalis (BA 47). This region overlaps with the lateral part of the orbitofrontal cortex (OFC) (Mackey & Petrides, 2010; Öngür, Ferry, & Price, 2003; Uylings et al., 2010) and anterior part of the ventrolateral prefrontal cortex (VLPFC), which have been implicated in a wide range of learning and decision-making behaviors, including the learning, maintenance and shifting of higher-order rules (Brass & von Cramon, 2004; Hampshire & Owen, 2006; Rushworth et al., 2005; Wallis, Anderson, & Miller, 2001; Wallis, Dias, Robbins, & Roberts, 2001), reversal learning (Kim & Ragozzino, 2005), extinction learning (Butter, Mishkin, & Rosvold, 1963), and value-based decision-making (see Wallis, 2011, for a review). Recently, Wilson, Takahashi, Schoenbaum, & Niv (2014) have proposed that many of these functions could be unified under a model in which the orbitofrontal cortex (particularly lateral OFC) computes an animal’s location in a cognitive map of task space, akin to a state representation for reinforcement learning. Such representations are particularly important when abstract information in working memory is necessary to distinguish between perceptually similar, but conceptually different, task states. Thus, it is possible that pattern change in the right pars orbitalis reflects changes in task context, which are difficult to infer from the physical environmental alone. Although our participants were not instructed to perform any task while encoding the story, understanding the narrative requires inferring the goals and situation of the characters, which is similar to navigating a complex task space. This interpretation is supported by studies showing that people understand stories by simulating the characters’ actions and that such simulation engages the same circuits involved in performing those actions in the real world (Speer, Reynolds, Swallow, & Zacks, 2009).

Extending our anatomical ROI analysis to the entire brain showed that, in addition to the right entorhinal and pars orbtalis, pattern change in the left caudal anterior cingulate cortex (ACC) predicted subsequent duration estimates. However, caudal ACC exhibited more rapid pattern change than the entorhinal and orbitofrontal cortex, and pattern change in this region did not correlate with the other two regions nearly as strongly as they did with each other, suggesting that it may represent a qualitatively different, faster-changing signal. Caudal ACC activity has been shown to increase in response to shifts in task contingencies (see Shenhav, Botvinick, & Cohen, 2013, for a review) and there is converging evidence that ACC responses are important for adjusting behavior to unexpected changes by increasing attention and learning rate (Bryden, Johnson, Tobia, Kashtelyan, & Roesch, 2011; Behrens, Woolrich, Walton, & Rushworth, 2007; McGuire, Nassar, Gold, & Kable, 2014). Furthermore, O’Reilly et al. (2013) have provided evidence that the ACC only responds to surprising outcomes when they necessitate updating beliefs about the current state of the world. Although the present study was not designed to test such accounts, our findings are consistent with a role for ACC in updating predictive models. Events in the story that prompt participants to update their beliefs about the characters’ situation are also likely to cause changes in cognitive context and therefore overestimation of duration. However, future studies are needed to test this interpretation, for instance by manipulating belief updating independently of surprise and measuring its effect on retrospective duration estimates. In addition to the anatomical ROI analysis, we performed a whole-brain searchlight that yielded an extensive cluster covering the right anterior temporal lobe, extending from the medial temporal regions described above to the middle temporal gyrus and temporal pole. Prior work has suggested that the middle temporal gyrus and temporal pole are involved in narrative comprehension (Ferstl, Neumann, Bogler, & Von Cramon, 2008; Mar, 2004) and narrative item memory (Hasson, Nusbaum, & Small, 2007; Maguire, Frith, & Morris, 1999). Moreover, Ezzyat and Davachi (2011) found a similarly located cluster (extending from the right perirhinal cortex to the right middle temporal gyrus) to be involved in integrating information within narrative events. In particular, they showed that activity within these regions gradually increases within events and that this increase predicts the degree to which memories become clustered within events. Retrospective time judgments have been shown to increase with the number of events an interval contains (Poynter, 1983; Zakay et al., 1994), suggesting that brain regions involved in clustering memories by events may carry important information for estimating durations.

## Conclusion

After probing human participants’ time perception for intervals from an auditory story they had just heard, we found substantial variability in subjective estimates of the passage of time. This variability was significantly correlated with changes in BOLD activity patterns in the right entorhinal cortex, right pars orbitalis, left caudal ACC, and the right anterior temporal lobe, between the start and end of each interval. Control experiments demonstrated that duration estimates were strongly driven by contextual boundaries and that the relationship between neural distance and behavior could not be explained by lower-level changes in story content. Our findings suggest that patterns of activity in these regions might encode contextual information that participants can later retrieve to infer the durations of intervals on the scale of minutes. Additional work is needed to assess how these regions contribute to representing particular contextual features (such as physical environment, abstract task states, and emotional states) and whether changes in each of these features affect retrospective duration estimates differently.

## Methods

### Participants

18 participants (13 female) took part in the study. All participants were recruited from the Princeton undergraduate and graduate student population and were between 18 and 31 years of age (mean = 22 years). All participants were screened to ensure no neurological or psychiatric disorders. Written informed consent was obtained for all participants in accordance with the Princeton Institutional Review Board regulations. Participants were compensated $20/hour for the scanning session, and $12/hour for the behavioral session.

### Experimental Design and Stimuli

The experiment consisted of two parts: an approximately 40-minute session in the MRI scanner, during which participants listened to the auditory story, followed immediately by a 1-hour behavioral session, during which participants completed a time perception test on the story they had just heard. ***Figure 1*** illustrates the experimental procedure.

### fMRI session

Prior to the fMRI session, participants were instructed to listen carefully to the auditory story while in the scanner, because they might be asked questions about it later. The nature of the follow-up questions was unknown to the participants. While in the scanner, participants listened to a 25-minute-long radio adaptation of a science fiction story called “Tunnel Under the World” (written by Frederik Pohl), originally aired on the radio drama series, “X Minus One”, in 1956.

### Time perception test

After leaving the scanner, participants were surprised with a time perception test, presented on a laptop with the Psychophysics toolbox for Matlab (Brainard, 1997; Pelli, 1997). For each of 43 questions, participants listened to a 10 s clip from the story, followed by another 10 s clip, and were asked to estimate how much time had passed between the first and second clips when they initially heard the story. Participants were specifically asked to estimate how much time had passed in their own lives, rather than how much narrative time had passed in the story. They were also asked to make the judgments as intuitively as possible, without resorting to deductive reasoning about the sequence of events that unfolded in between the two excerpts.

Participants had complete control over the pacing of the test. On each question, they initiated the playing of the clips, and were able to replay the clips if they missed them the first time. They could take as long as they wished to enter their duration estimates (in minutes and seconds), using the keyboard. Clip pairs were identical across participants, but the order in which the pairs were presented was randomized.

To control for the objective passage of time, we ensured that 24 of the clip pairs were 2 minutes apart and 19 of the pairs were 6 minutes apart. Debriefing showed that participants were unaware of this manipulation, and the high variability of duration estimates for both the 2 and 6-minute intervals further confirmed that they were unaware of the fixed interval durations.

After participants had provided duration estimates for all 43 intervals, the 86 clips that had delimited those intervals were replayed in a random order (unpaired), and participants were asked to place each clip on the timeline of the story. For each of the 86 questions, a white line appeared on a black background, representing the full length of the story. Participants could place their cursor at any point on that line, followed by the Enter key. After each placement, they were asked to provide a confidence rating on a scale of 1 to 5, reflecting their confidence about that clip’s place in the story. Participants were instructed to base the confidence rating on their certainty of when that clip occurred in the story, rather than on the vividness of the memory for that clip. While the exact placement of each clip on the timeline was not used in the fMRI analysis, confidence ratings were used to exclude clips whose temporal context participants had forgotten.

Please note: the first of our 18 participants completed a version of the time perception test that differed only in the following way: the specific intervals in the story whose duration was asked about were different. In all other respects (half of the intervals were 2 minutes while the other half were 6 minutes apart), the behavioral test was identical to the subsequent 17 participants. For this reason, however, any analyses where duration estimates are compared across participants were performed on 17 rather than 18 participants. Any within-participant analyses were performed on all 18 data sets.

### Naïve time perception test

To address the concern that participants were estimating temporal distance between two clips based purely on the content of the clips (rather than their memory of when the clips had occurred in the story), we administered an identical time perception test to a separate group of 17 participants who had never heard the story. Since these participants had no memory of the story, they could only base their temporal distance estimates on the content of the clips. Participants were only told the length of the story (25 minutes, 33 seconds) and informed that the distance between two clips could not be greater than this length.

### Event boundary test

A separate group of 9 participants were asked to listen to the same story and to press the space bar every time they thought an event had ended and a new event was beginning. This test was purely behavioral and fMRI data were not collected for these participants.

### MRI Acquisition

Participants were scanned in a 3T full-body MRI scanner (Skyra, Siemens) with a 20-channel head coil. Functional images were acquired using a T2*-weighted echo planer imaging (EPI) pulse sequence (repetition time [TR], 1500 ms; echo time [TE], 28 ms; flip angle, 64°), each volume comprising 27 slices of 4 mm thickness. In-plane resolution was 3×3 mm^2^ (field of view [FOV], 192×192 mm^2^). Slice acquisition order was interleaved. Anatomical images were acquired using a T1-weighted magnetization-prepared rapid-acquisition gradient echo (*M*PRAGE) pulse sequence (TR, 2300 ms; TE, 3.08 ms; flip angle 9°; 0.89 mm^3^ resolution; FOV, 256 mm^2^). Participants’ heads were stabilized with foam padding to minimize head movement. Auditory stimuli were presented using the Psychophysics toolbox (Brainard, 1997; Pelli, 1997). Participants were provided with MRI compatible in-ear mono earbuds (Sensimetrics Model S14), which provided the same audio input to each ear. MRI-safe passive noise-canceling headphones were placed over the earbuds for additional protection against noise.

### fMRI Data Preprocessing

FMRI data processing was carried out using FEAT (FMRI Expert Analysis Tool) Version 5.98, part of FSL (FMRIB’s Software Library, www.fmrib.ox.ac.uk/fsl). The following procedure was applied: motion correction using MCFLIRT (Jenkinson, Bannister, Brady, & Smith, 2002); slice-timing correction using Fourier-space time-series phase-shifting; non-brain removal using BET (Smith, 2002); spatial smoothing using a Gaussian kernel of FWHM 6.0 mm; grand-mean intensity normalization of the entire 4D dataset by a single multiplicative factor; and high-pass temporal filtering (Gaussian-weighted least-squares straight line fitting, with sigma=240.0s). The procedure for selecting the high-pass filter is described below. Preprocessed data were kept in the native functional space for all analyses.

### Procedure for obtaining anatomical masks: FreeSurfer and MTL segmentation

Segmentation was performed in a semi-automated fashion using the FreeSurfer image analysis suite, which is documented and available online (version 5.1; http://surfer.nmr.mgh.harvard.edu) with details described previously (e.g. Fischl et al., 2004; Poppenk & Norman, 2014). Briefly, this processing includes removal of non-brain tissue using a hybrid watershed/surface deformation procedure (Ségonne et al., 2004), automated Talairach transformation, intensity normalization (Sled, Zijdenbos, & Evans, 1998), tessellation of the grey matter / white matter boundary, automated topology correction (Fischl, Liu, & Dale, 2001; Segonne, Pacheco, & Fischl, 2007), surface deformation following intensity gradients (Fischl & Dale, 2000), parcellation of cortex into units based on gyral and sulcal structure (Desikan et al., 2006; Fischl et al., 2004), and creation of a variety of surface-based data, including maps of curvature and sulcal depth.

We resampled and aligned FreeSurfer segmentations of all grey matter, white matter, and cerebrospinal fluid (CSF) regions to native functional image space for use as anatomical masks. Anatomical regions were segmented according to the Desikan-Killiany Atlas (Desikan et al., 2006).

It is important to note that the medial temporal lobe (MTL) masks in the Desikan-Killiany Atlas do not match the canonical anatomical distinctions in the literature. For example, the parahippocampal gyrus mask comprises the medial part of the parahippocampal cortex and the posterior part of the entorhinal cortex. Therefore, instead of the FreeSurfer MTL masks, we used a probabilistic MTL atlas developed by Hindy & Turk-Browne (2015). MTL regions, including perirhinal cortex, entorhinal cortex and parahippocampal cortex were defined probabilistically in MNI space, based on a database of manual MTL segmentations from a separate set of 24 participants. Manual segmentations were created on *T*_2_-weighted turbo spin-echo images using anatomical landmarks (Duvernoy, 2005; Carr, Rissman, & Wagner, 2010; Schapiro, Kustner, & Turk-Browne, 2012), and then registered to an MNI template. Finally, nonlinear registration (FNIRT; Andersson, Jenkinson, & Smith, 2007) was used to register the masks from MNI space to each participant’s native space. After registration, voxels with a probability greater than 0.3 of being in a region were assigned to that ROI.

### Residualization of non-neuronal signal sources

Slow changes of respiration over time (RV) have been shown to induce robust changes in the BOLD signal (Chang, Cunningham, & Glover, 2009) in many areas around the cerebral midline. To minimize signal change unrelated to neural activity, we used multiple linear regression to project out 3 nuisance variables from the BOLD data (Behzadi, Restom, Liau, & Liu, 2007; Silbert, Honey, Simony, Poeppel, & Hasson, 2014). Nuisance regressors were:

1) the average time course of high standard deviation voxels (voxels with the top 1% largest standard deviation), as these voxels tend to have the highest fractional variance of physiological noise (e.g., cardiac and respiratory components) and are likely near blood vessels (Behzadi et al., 2007),
2) the average BOLD signal measured in CSF,
3) the average white matter signal.

All masks (grey matter, white matter and CSF) were obtained from the FreeSurfer segmentation procedure described above. The beneficial effects of this residualization procedure on the signal-to-noise ratio are shown in ***Figure 10***. Note that this procedure was always applied after removal of low-frequency components using the high-pass filter (see below.)

**Figure 10.**
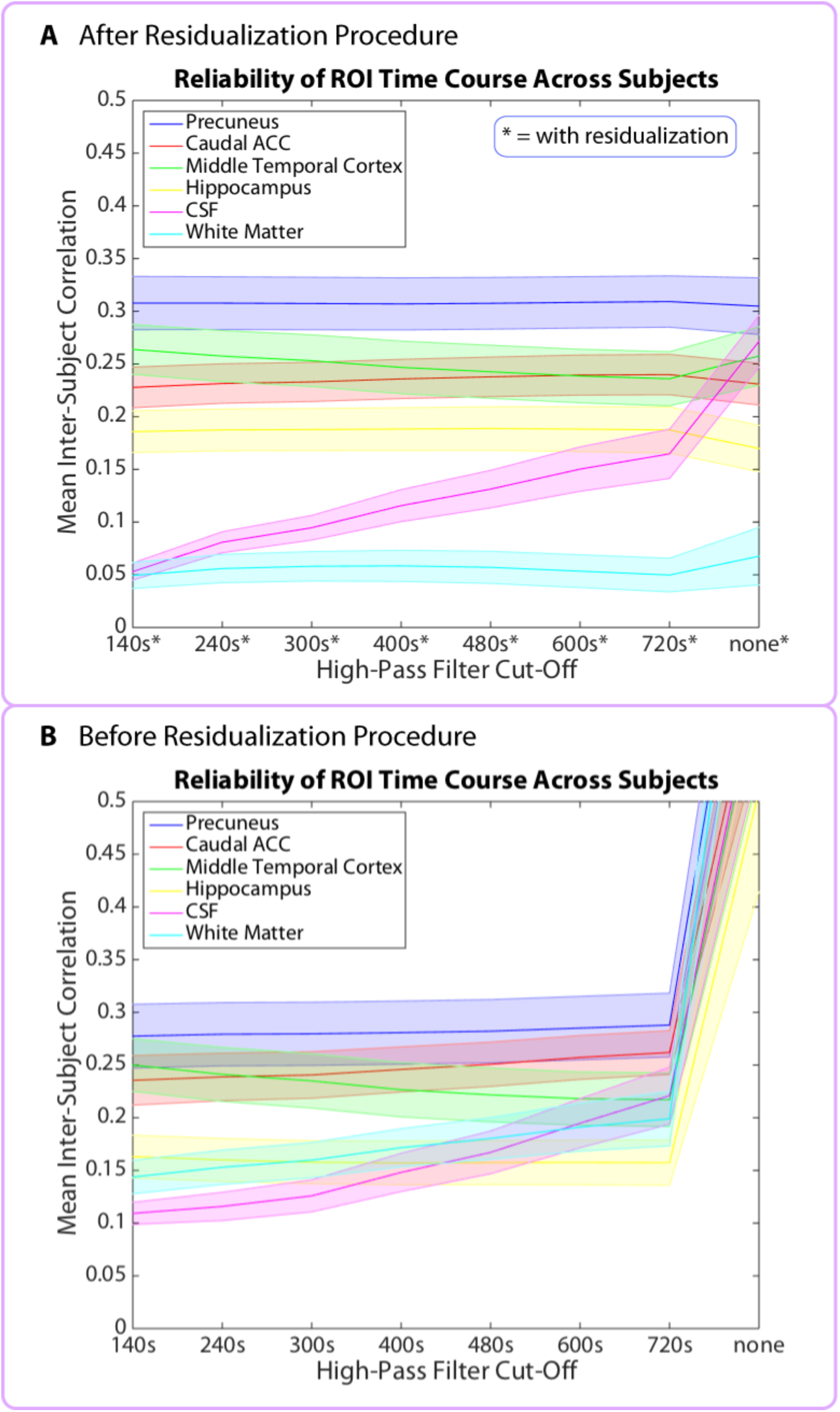
Mean inter-subject correlations (ISCs) for 6 representative brain regions as a function of the high-pass filter cut-off. Shaded error bars represent standard errors of the mean (across participants). Top panel **A** shows the mean ISCs after the residualization procedure has been applied (see “Residualization of non-neuronal signal sources”). The 480 s cut-off was the gentlest filter for which all of the grey matter regions listed above showed ISC values significantly above those in the CSF. Bottom panel **B** shows the mean ISCs prior to the residualization procedure. Without residualization, the ISCs of some grey matter regions never rise significantly above those in the white matter and CSF. Note that without high-pass filtering (“none”) or residualization, all brain regions displayed spuriously high ISCs.

### Methodological challenges with analyzing pattern distance over long time scales: Selection of temporal high-pass filter cut-off

Because we were interested in the aspect of neural activity that changes slowly over time (reflecting gradual changes in context), we could not use a standard high-pass filter (with a cut-off period on the order of 120 s), as it would remove components of the signal that evolve on the scale of minutes. Thus, we were faced with the challenge of preserving slower components of the BOLD signal that reflect neural activity, while removing low-frequency components attributable to non-neuronal noise, including scanner drift and physiological noise (such as low-frequency respiratory variation and heart rate variation). Physiological noise (and a substantial component of scanner noise) was factored out using the residualization procedure described above. This enabled us to select a gentler high-pass filter than is generally used in the literature.

We then performed a separate analysis to determine the optimal high-pass filter cut-off period, i.e. the lowest frequency cut-off that still enabled us to remove most of the non-neuronal noise. This analysis relies on the idea that, when participants listen to the same story or watch the same film, the signal in brain regions processing the story is highly correlated across participants (Hasson, Nir, Levy, Fuhrmann, & Malach, 2004). While such correlations should not be present in CSF or white matter, spurious inter-subject correlations in these regions can arise due to low-frequency noise. In addition, listening to the same story could induce correlated motion across participants, but these correlations would also be present in CSF and white matter. Thus, we searched for a high-pass filter that could remove nonspecific correlations in CSF and white matter, while preserving correlations in brain regions known to be important for processing the stimulus. For each participant, the inter-subject correlation (ISC) of a brain region was defined as the correlation between that participant’s ROI time course (averaged over voxels in that region) with the average time course of all the other participants (Hasson, Yang, Vallines, Heeger, & Rubin, 2008; Lerner et al., 2011).

Since the functional scan length was 1560 s (26 minutes), high-pass filter cut-off periods of 140 s, 240 s, 300 s, 400 s, 480 s, 600 s and 720 s were attempted. The minimal cut-off attempted, 140 s, was the cut-off used in several previous studies with naturalistic stimuli (e.g. Lerner et al., 2011), while 720 s represented approximately half of the scan duration and was the longest cut-off that could reasonably make a difference to data quality.

Given that roughly half the clip pairs in our time perception test were 2 minutes apart and the other half were 6 minutes apart, we hoped to find a filter that would allow us to measure pattern distances at both of these time scales. However, we were unable to find a high-pass filter that would allow us to examine activity patterns that were 6 minutes (360 s) apart. In order to meaningfully measure distances between neural patterns that are 360 s apart, the Nyquist theorem suggests we would need a high-pass filter cut-off of 720 s or larger. However, plotting ISC as a function of high-pass filter (***Figure 10***) showed that a cut-off like 720 s was not able to remove inter-subject correlations in the CSF, which remained of the same magnitude as those in some grey matter regions. We concluded that pattern distances at the 6-minute time scale are too confounded with low-frequency noise (as reflected in spurious correlations in the CSF), and therefore restricted our analysis to intervals that were 2 minutes long.

According to the Nyquist theorem, we need a filter cut-off of 4 minutes (240 s) or longer in order to measure distances between patterns that are 2 minutes apart (120 s). Out of the filters tested (240 s – 720 s), a cut-off of 480 s was selected to be the gentlest (i.e. the longest) filter that reduced the magnitude of inter-subject correlations in ventricles and CSF, such that they were significantly below the correlations in most grey matter regions.

***Figure 10*** illustrates that, even for regions like the hippocampus – with relatively low inter-subject correlations – the 480 s filter cut-off, combined with the residualization procedure, succeeded at raising the grey matter ISCs significantly above those of the white matter and CSF.

### fMRI Data Analysis

#### Within-participant correlation between pattern change and duration estimates

Our primary hypothesis was that greater pattern dissimilarity between two clips (at the time of encoding) would correlate with greater subsequent duration estimates. For each pair of clips from the time perception test, we located the TRs (volumes) corresponding to when the participant first heard those clips and extracted the activity patterns for each ROI at those time points. Since the auditory clips were between 5 s and 10 s in duration (corresponding to about 5 volumes), we averaged the patterns over 5 consecutive TRs for every clip, with the 5-TR window centered on the middle of each clip.

We then related the pattern distance between the two clips at encoding to how much time the participant thought passed between them. Specifically, we calculated the dissimilarity (1 – Pearson correlation) between the two averaged activity patterns. The pattern dissimilarity scores for a given region were then correlated with that participant’s subsequent duration estimates. This was performed separately for every ROI and searchlight (***Figure 3***). We thus obtained a Pearson correlation score for every ROI in every participant. To assess the reliability of the correlation across participants for a given ROI, we ran a phase-randomization procedure, which is described in detail below. The results of the phase-randomization procedure were then subjected to multiple comparisons correction.

#### Removing low-confidence intervals

If a participant could not remember when in the story a particular clip had occurred, it would be difficult for them to estimate the temporal distance between that clip and another clip. It is possible that participants would invoke different retrieval strategies in such cases (for instance, they might base their duration estimates purely on the content of the clips, without recollecting their context). It is also possible that such estimates could be random guesses. To filter out guesses, we used the confidence ratings collected after the time perception test, in which participants rated how well they could remember when in the story each individual clip had occurred. Specifically, we located the participant’s confidence for the two clips delimiting each temporal interval, and took the smaller of the two ratings as the confidence for that interval. We performed the main analysis relating neural drift to time estimation (described below) only on high-confidence intervals, removing pairs of clips with the lowest confidence. Since participants calibrated their confidence ratings differently (some were more prone to rate their confidence as 4/5, while others were more prone to rate it as 2/5), we picked the confidence threshold for each participant that removed at least 33% of the intervals with the lowest confidence, while preserving at least 33% of the intervals with the highest confidence. Our behavioral analysis (see *Results*) shows that participants’ duration estimates were significantly more accurate for high-confidence intervals than when all intervals were included.

#### Statistical analysis of correlations between pattern change and behavior

Because of the presence of long-range temporal autocorrelation in the BOLD signal (Zarahn et al., 1997), the statistical likelihood of each observed correlation (between neural distance and duration estimates) was assessed using a permutation procedure based on surrogate data. The surrogate data were generated using phase randomization (Theiler et al., 1992). Phase-randomized surrogates have the same autocorrelation as the original signal.

Since our analysis measures pattern change over multiple voxels, rather than the time course of a single voxel, we generated surrogate time courses of pattern change (***Figure 3 – Supplement 1*** shows how that time course was obtained). Having extracted the time course of pattern change for each ROI, we applied a Fourier transform to that signal. To randomize its phases, we multiplied each complex amplitude by *e*^*jφ*^, where ^*φ*^ is independently chosen for each frequency from the interval [0, 2π]. In order for the inverse Fourier transform to be real (no imaginary components), we symmetrized the phases, so that ^*φ*(*f*) = ‒*φ*(–*f*)^. Finally, we took the inverse Fourier transform to produce the surrogate time courses.

Each surrogate dataset was analyzed in the same manner as the empirical data: pattern dissimilarity between each pair of clips was correlated with duration estimates. Thus, we generated a distribution of 10,000 null correlations for every ROI in every participant (see ***Figure 3 – Supplement 1***). For every ROI, we were then able to compare the empirical Pearson correlation with the distribution of null correlations. We calculated a Z-value for every participant:

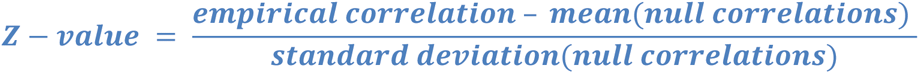

A large positive Z-value implies that the empirical correlation is large relative to the distribution of null correlations. To assess whether the Z-values for a given ROI were reliably positive across participants, we performed a right-tailed t-test against 0. The p-values from the above t-test were then subjected to multiple comparisons correction. For anatomical ROIs (derived from the FreeSurfer and MTL atlases), we used False Discovery Rate (FDR) to correct for multiple comparisons. This procedure returns q-values (FDR adjusted p-values) using the linear-step up (LSU) procedure introduced by Benjamini & Hochberg (1995). For the searchlight analysis, we controlled the family-wise error (FWE) rate, as described below.

### ROI selection

The literature reviewed above suggested that the MTL, lateral prefrontal cortex, insula, putamen and inferior parietal cortex might all process information important for inferring the duration of past events. We therefore performed an ROI analysis on the following regions, derived from both the FreeSurfer and MTL atlases: hippocampus, parahippocampal cortex, entorhinal cortex, perirhinal cortex, amygdala, superior frontal cortex, caudal and rostral middle frontal gyrus (dorsolateral prefrontal cortex), pars opercularis (frontal operculum), pars triangularis, pars orbitalis, lateral orbitofrontal cortex, frontal pole, insula, putamen and inferior parietal cortex. This resulted in an analysis on 16 regions of interest (in each hemisphere) motivated by the literature. ROIs with q-values < 0.05 (FDR) are reported as significant.

As part of an exploratory, whole-brain search, we also ran the same analysis on all grey matter regions in the Desikan-Killiany Atlas, which contained 42 regions in each hemisphere, including the ones mentioned above (see *FreeSurfer Segmentation and MTL Segmentation*). The complete list of regions can be found in ***Figure 4 – Supplement 1***. For the exploratory analysis, we report regions with q-values < 0.1 (FDR).

### Whole-brain searchlight

In addition to using anatomical ROIs, we ran a cubic searchlight with 3x3x3 (27) voxels throughout the entire brain. The same analysis as described above was performed for every searchlight, and the Z-value for each searchlight was assigned to the center voxel. Each participant’s Z-value map was then transformed to standard MNI space and down-sampled to 3mm to reflect the resolution of the original data. Family-wise error rate was controlled using FSL’s *randomise* function (version 5.0.4, Winkler, Ridgway, Webster, Smith, & Nichols, 2014). An uncorrected p-value image was first generated, reflecting voxel-wise (searchlight) reliability across participants. The significance of supra-threshold clusters (defined by the cluster-forming threshold, *p*<0.01) was then assessed by cluster mass. Specifically, a corrected p-value was assigned to each cluster by assessing its cluster mass with respect to the null distribution of the maximum cluster mass during 10,000 permutation simulations (Hayasaka & Nichols, 2003; Nichols & Holmes, 2002). Cluster coordinates are reported in MNI space, and cluster size reflects the number of voxels in 3x3x3mm MNI space.

### Within-interval correlation between pattern change and duration estimates

Our main analysis verified whether the pattern distance between two clips was correlated with duration estimates in a given participant and then aggregated the results across participants. To address the concern that pattern distance between two clips might reflect only the difference in story content between those clips (rather than change in abstract factors like mental context), we performed the same analysis for a given interval across participants and aggregated the results across intervals. Since this analysis is performed within intervals, it ensures that story content is held constant across participants, such that differences in pattern distances and duration estimates are due to individual differences only. To ensure that pattern distances and duration estimates were comparable across participants, all vectors were z-scored within participant. The Pearson correlation between pattern distances and duration estimates across participants was then calculated for every 2-minute interval in every ROI.

As for the main analysis, this analysis was performed on high-confidence intervals. For each interval, we only included participants who had confidently recollected the temporal position of the two clips delimiting that particular interval.

The significance of each correlation score was assessed using a permutation test: 10 000 null correlations were obtained by scrambling the duration estimates across participants, such that a given participant’s duration estimate was matched with a different participant’s pattern distance. Since this analysis was performed across participants, it was not necessary to generate phase-randomized pattern distance time courses – the auto-correlation in the BOLD signal for a given participant only plays a role for the within-participant analysis.

As above, a Z-value was obtained for every interval, reflecting the degree to which the empirical correlation was higher than the distribution of null correlations. Finally, a right-tailed t-test was performed to assess whether a given ROI’s Z-values were reliably positive across intervals. The p-values from this t-test were subjected to multiple comparisons correction using FDR.

To compare effect sizes between the within-interval and within-participants analyses, we calculated Cohen’s *d* for a region as:

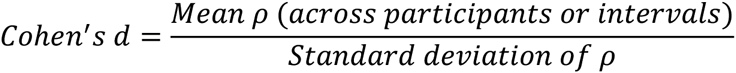

where *ρ* is the Pearson’s correlation between pattern distance and duration estimates.

(Using the Z-values derived from the permutation procedures, rather than the raw correlation scores, yielded practically identical results.)

#### Comparing speed of pattern change across brain regions

If the brain regions found significant in our main analysis represent mental context, then the pattern of activity in these regions should change more slowly over time than the patterns in regions representing sensory information. To quantify the speed of pattern change in a given ROI, we obtained the correlation of the pattern at every time point (TR) with itself at every other time point. (As for our main analysis, the BOLD time course of every voxel was smoothed using a moving average filter of 5 TRs. This temporal smoothing was used as a de-noising technique and did not affect the results.) We then averaged the auto-correlation curves across TRs to obtain a mean auto-correlation function for every region in every participant. The more rapidly a pattern changes over time, the more sharply the auto-correlation should decrease as we move away from 0. To quantify this, we defined the Full-Width Half-Maximum (FWHM) of the auto-correlation curve as the number of time points (TRs) for which the auto-correlation was equal to or greater than half its maximum value (the maximum was always 1.)

To compare the speed of pattern change in the regions we found (pars orbitalis, right entorhinal cortex and left caudal ACC) with regions involved in auditory and language processing (right transverse temporal cortex, which contains primary auditory cortex, as well as the right superior temporal cortex and right banks of the superior temporal sulcus), we performed a paired Wilcoxon signed rank test on the FWHM values across participants. The p-values from this test were subjected to multiple comparisons correction using FDR.

Since the anatomical masks we used varied substantially in size, we sought to ensure that differences in the speed of pattern change were not due to differences in ROI size. For this purpose, we performed the same analysis after regressing the vector of ROI sizes out of the vector of FWHM values for every participant.

Since the above regression would only account for a linear effect of ROI size on the speed of pattern change, we additionally performed a univariate analysis that calculated the auto-correlation function for each voxel individually. The auto-correlation curve was obtained by correlating the BOLD time course of every voxel with itself at all possible lags. The mean auto-correlation for an ROI was obtained by averaging the auto-correlation curves across all the voxels in that ROI. The FWHM values were then calculated in the same manner as above for every ROI in every participant.

### Behavioral Data Analysis

#### Significance of correlation between duration estimates and event boundaries

To assess whether the number of event boundaries in an interval predicted duration estimates for that interval, we related our original participants’ duration estimates with event boundary data collected from a separate group of 9 participants. For each 2-minute interval from the time perception test, we counted the number of event boundaries that a participant had indicated during that interval and averaged that number across the 9 participants. This resulted in a mean number of event boundaries per interval, which was then correlated with the mean estimated duration of that interval from our original participants.

To assess the statistical significance of this correlation, we performed a bootstrapping procedure on the duration estimates. We obtained 1000 bootstrap samples, each time selecting with replacement a different subset of *n* individuals from our pool of *n* participants. The duration estimates for each subset were averaged across participants and correlated with the mean number of event boundaries. The upper limit (*ul*) for an *×%* confidence interval was set to the value of the Pearson correlation in percentile *×%* of the bootstrap distribution; the lower limit (*ll*) for the confidence interval was set to the value of the beta score in percentile 100-*×* of this distribution. Confidence intervals that did not encompass zero were considered reliable at the given level of confidence.

#### Significance of difference in correlations with event boundaries between original duration estimates and naïve duration estimates

We hypothesized that duration estimates from our original participants (who had actually heard the story) would be significantly more correlated with the number of event boundaries between two clips than duration estimates from our naïve participants, who had never heard the story. To assess the significance of the difference in correlations, we computed the *r*_*diff*_ (empirical difference), as well as the upper confidence limits (*ul*_*diff*_) and lower confidence limits (*ll*_*diff*_) for the difference between the two correlations. We used the following formulae (Zou, 2007; Poppenk & Norman, 2012) for two bootstrapped correlation confidence intervals:

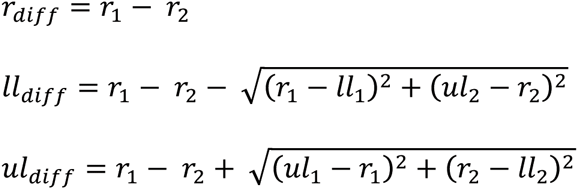

The upper (*ul*_1_, *ul*_2_) and lower limits (*ll*_1_, *ll*_2_) for a 95% confidence interval of each group’s correlation were calculated as described above.

#### Reliability of duration estimates across participants within and between groups

We hypothesized that both our original participants and the naïve participants (who had never heard the story) would use consistent strategies to estimate the temporal distance between two clips, but that these strategies would differ across groups. If this is the case, duration estimates should be more reliable across participants within groups than across participants between groups.

To assess within-group reliability, we correlated each participant’s duration estimates with the mean of the other participants’ estimates. These correlations were then averaged across participants within a group to obtain a mean within-group ISC (inter-subject correlation). The between-group reliability was calculated by correlating each participant’s duration estimates from one group (e.g., the original participants) with the mean duration estimates from the other group (e.g., the naïve participants). These correlations were then also averaged across participants to obtain a mean between-group ISC.

To assess the significance of the difference between the mean within-group ISC and the mean between-group ISC, we compared the empirical difference with a null distribution of differences. Group labels (naïve participants vs. original participants) were scrambled 10,000 times, such that each participant’s duration estimates were randomly assigned to either the naïve group or to the original group. The difference between the mean within-group ISC and the mean between-group ISC was then computed for these two random groups. Using this null distribution of ISC differences, we calculated a p-value based on the number of permutations that yielded a greater difference than the empirical difference.

Please note that the within-group and between-group correlations could be compared only because the group sizes were identical (17 participants in each) and because the within-group correlations were equally strong for the original and naïve groups (*M*=0.43, *SD*=0.25 vs. *M*=0.39, *SD*=0.24; *t*(32)=0.50, *p*=0.62). Since the within-group ISCs are comparable, we can infer that the significant difference between the within-group and between-group reliability reflects a difference in the signals (strategies) underlying the two groups of duration estimates (Chow, Chen, & Hasson, 2015), rather than a difference in within-group reliability.

## Acknowledgments

We would like to thank Lucy Lin for her assistance with data collection for the event boundary experiment. We would like to thank Erez Simony, Lili Sahakyan, Mariam Aly, Anna Schapiro and Michael Chow for their advice on data analysis and preprocessing, as well as helpful discussion.

---

**Figure 2 – Supplement 1:**
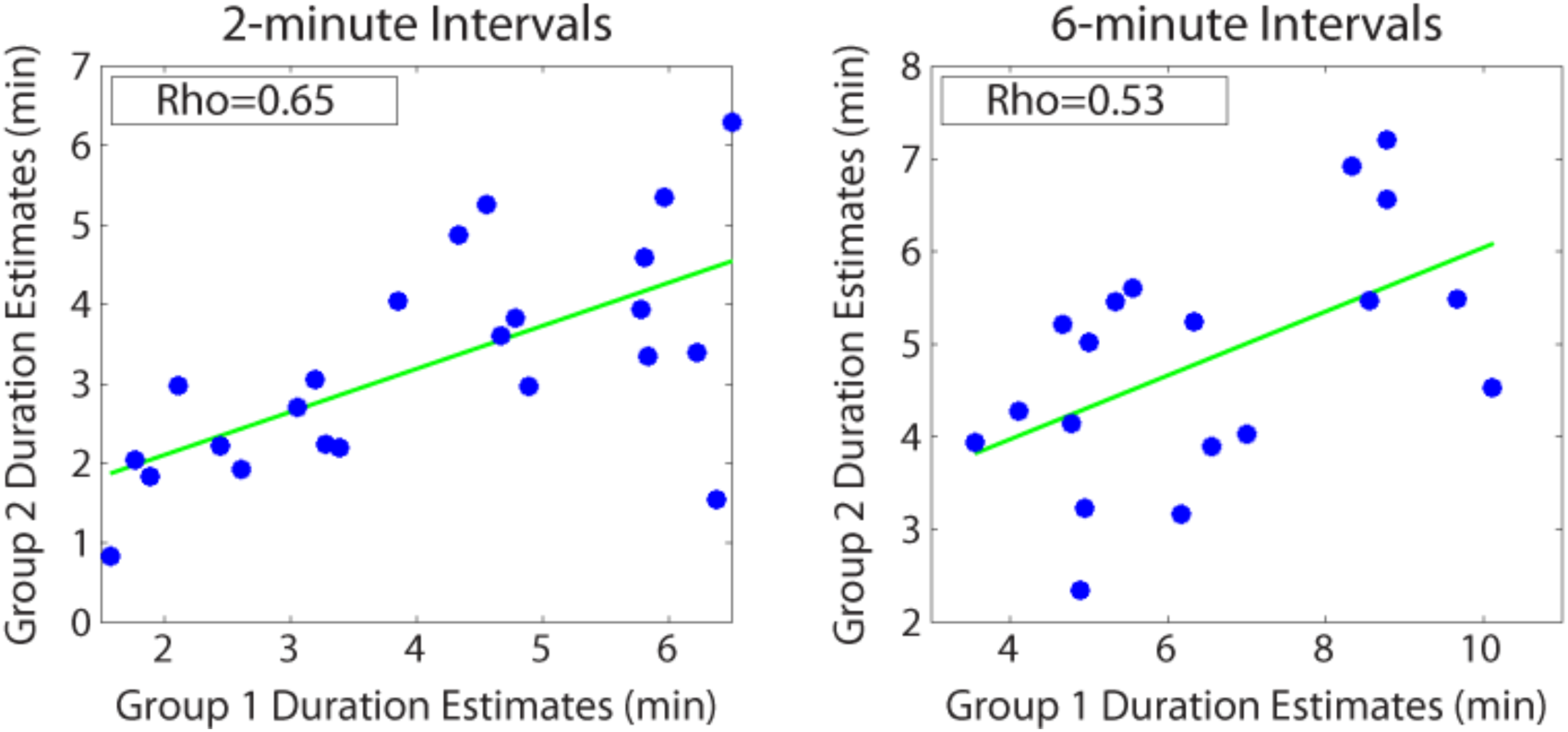
Reliability of duration estimates across participants. Between-group correlations were obtained by splitting the participants randomly into two equal groups and averaging the duration estimates for each interval (across participants) within a group. Each dot in the scatterplot represents a particular temporal interval; its *x* and *y* coordinates indicate the mean estimated duration of that interval for Group 1 and Group 2 participants, respectively. We repeated this procedure 1000 times to ensure that we sampled a variety of group splits. The average correlation between the two groups was 0.64 (*SD*=0.09) for 2-minute intervals and 0.54 (*SD*=0.15) for 6-minute intervals. The above plot shows the grouping that was most representative of the mean. This analysis suggests that features of the story made some intervals appear consistently shorter and other intervals appear consistently longer across participants.

**Figure 3 – Supplement 1:**
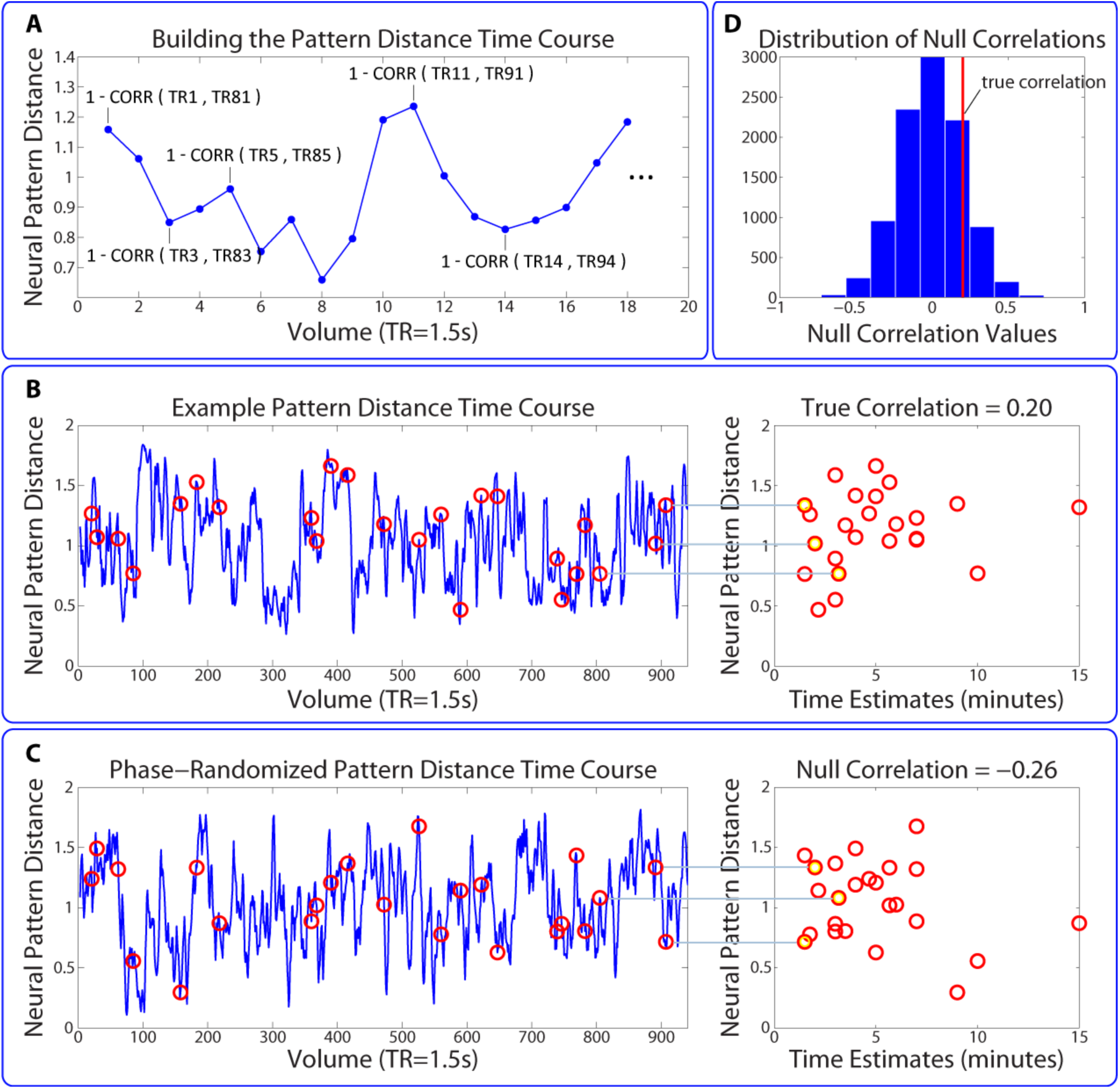
Permutation test assessing the temporal specificity of correlations between pattern change and behavior. This procedure is described in the *Methods* (see “Statistical analysis of correlations between pattern change and behavior”). (**A,B**) The time course of pattern change is constructed using the distance (1 - Pearson’s *r*) between each pattern and the pattern 80 TRs (2 minutes) after it. As in the main analysis, we averaged over the 5 consecutive TRs surrounding each pattern (for simplicity, this is not shown in the above figure). (**C**) 10 000 surrogate pattern distance time courses are generated by randomizing the phases of the original time course, thus conserving the amplitude of each frequency component. (**D**) Surrogate pattern distances are correlated with time estimates, generating 10,000 null correlations. A Z-value for each ROI / searchlight in each participant is computed to compare the strength of the empirical correlation with the distribution of null correlations. The p-value for a given ROI is obtained using a right-tailed t-test on the Z-values across participants.

**Figure 4 – Supplement 1:**
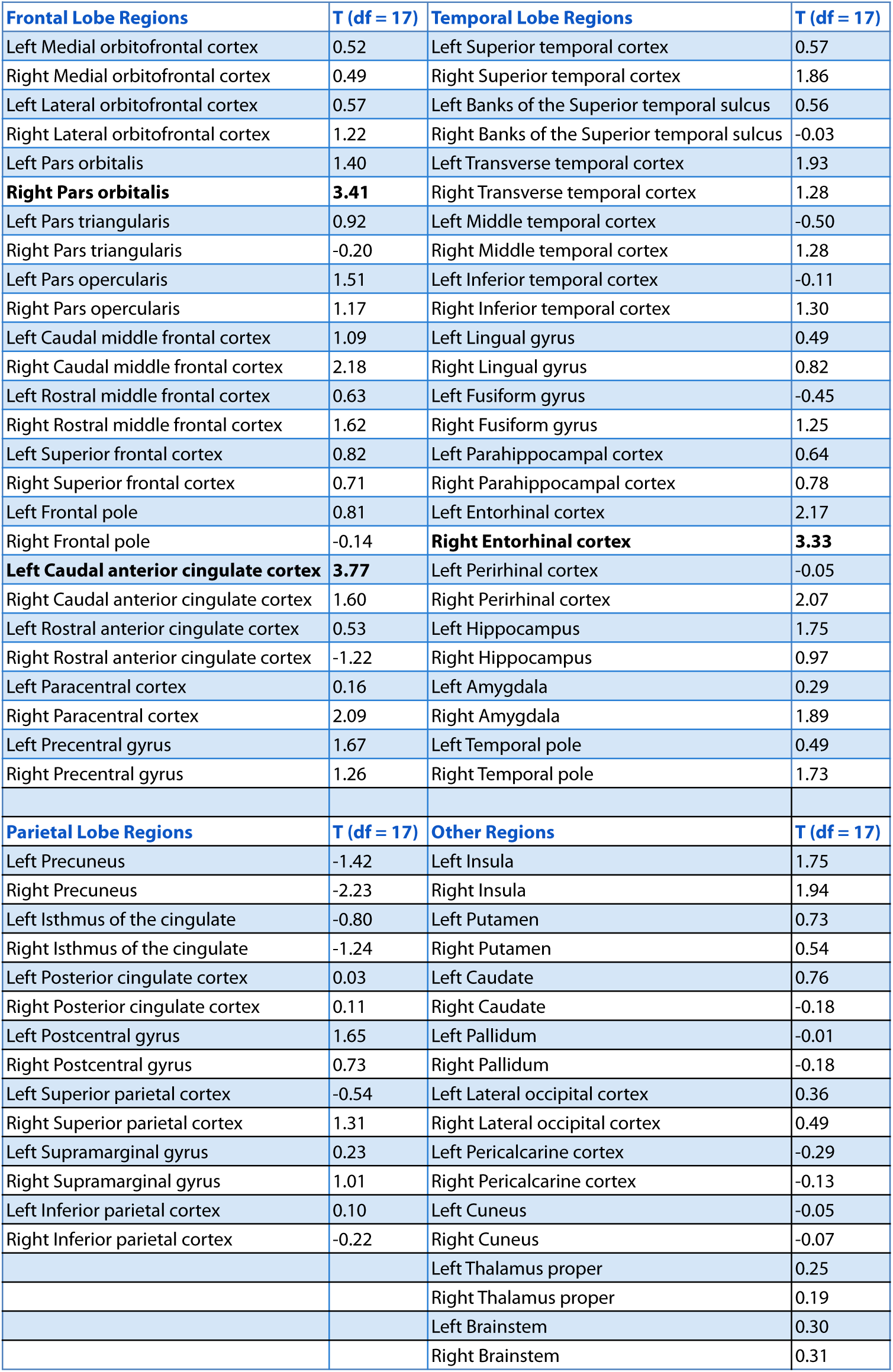
Whole-brain results for all grey matter regions derived from FreeSurfer segmentation and the probabilistic MTL atlas. T-values were obtained from a t-test verifying whether the Z-values for a region were reliably positive across participants (i.e., whether the empirical correlation between pattern distance and duration estimates was high relative to the distribution of null correlations). The three regions in bold survived whole-brain FDR correction at *q*<0.1.

**Figure 4 – Supplement 2:**
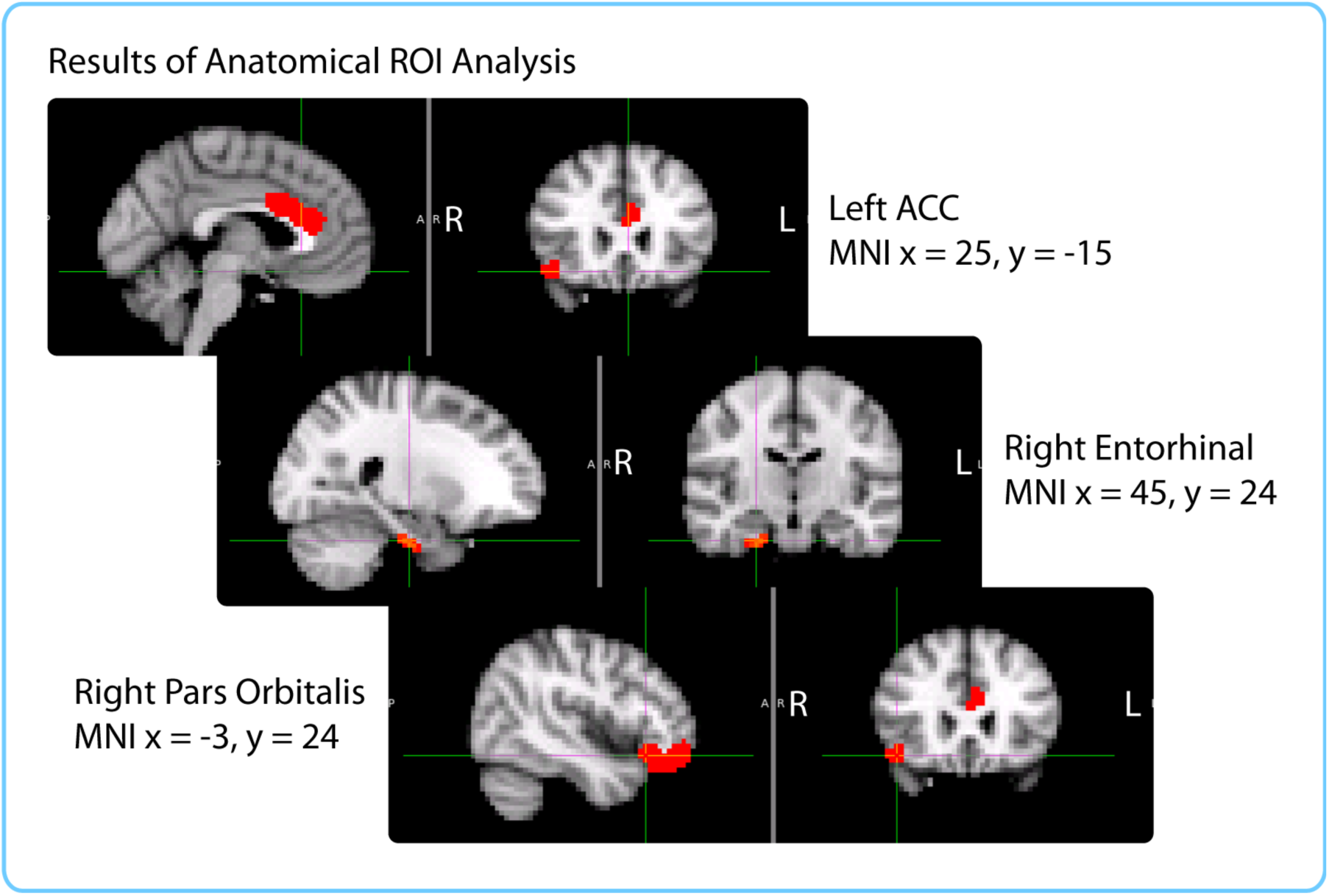
Anatomical ROIs that showed a significant correlation between pattern change and duration estimates within participants, after whole-brain FDR correction. In red are regions with q<0.1: the right entorhinal cortex, right pars orbitalis and left caudal ACC. This analysis was performed in native space on participant-specific ROIs. ROIs were transformed from native functional space to MNI space for display purposes.

